# Dormant replication origin firing links replication stress to whole chromosomal instability in human cancer

**DOI:** 10.1101/2021.10.11.463929

**Authors:** Ann-Kathrin Schmidt, Nicolas Böhly, Xiaoxiao Zhang, Benjamin O. Slusarenko, Magdalena Hennecke, Maik Kschischo, Holger Bastians

**Author notes:** these authors contributed equally to this work (Ann-Kathrin Schmidt, Nicolas Böhly, Xiaoxiao Zhang). corresponding author: Holger Bastians, Georg-August University Göttingen, Göttingen Center for Molecular Biosciences (GZMB) and University Medical Center Göttingen (UMG), Institute of Molecular Oncology, Section for Cellular Oncology, Grisebachstrasse 8, D-37077 Göttingen, Germany.

## Abstract

Chromosomal instability (CIN) is a hallmark of cancer and comprises structural CIN (S-CIN) and whole chromosome instability (W-CIN). Replication stress (RS), a condition of slowed or stalled DNA replication during S phase, has been linked to S-CIN, whereas defects in mitosis leading to chromosome missegregation and aneuploidy can account for W-CIN. It is well established that RS can activate additional replication origin firing that is considered as a rescue mechanism to suppress chromosomal instability in the presence of RS. In contrast, we show here that an increase in replication origin firing during S phase can contribute to W-CIN in human cancer cells. Increased origin firing can be specifically triggered by overexpression of origin firing genes including *GINS1* and *CDC45,* whose elevated expression significantly correlates with W-CIN in human cancer specimens. Moreover, endogenous mild RS present in cancer cells characterized by W-CIN or modulation of the origin firing regulating ATR-CDK1-RIF1 axis induces dormant origin firing, which is sufficient to trigger chromosome missegregation and W-CIN. Importantly, chromosome missegregation upon increased dormant origin firing is mediated by increased microtubule growth rates leading to the generation of lagging chromosomes in mitosis, a condition prevalent in chromosomally unstable cancer cells. Thus, our study identified increased or dormant replication origin firing as a hitherto unrecognized, but cancer-relevant trigger for chromosomal instability.

## Introduction

Chromosomal instability (CIN) is a hallmark of human cancer and correlates with tumor progression, development of therapy resistance, and poor clinical outcome ^1–3^. CIN can be categorized into two major forms: numerical or whole chromosomal instability (W-CIN) leading to aneuploidy and structural chromosomal instability (S-CIN), which causes structural chromosomal aberrations including deletions, insertions, and amplifications ^2^. S-CIN can be mechanistically traced back to errors in DNA repair and, in particular, by abnormal or slowed-down DNA replication, a condition known as replication stress (RS) ^4–6^. On the other hand, W-CIN is considered to be caused by errors during chromosome segregation in mitosis. In fact, various defects during mitosis have been suggested to contribute to W-CIN including supernumerary centrosomes, spindle abnormalities or impaired spindle checkpoint function ^1,7,8^. It is well established that a major mitotic abnormality in chromosomally unstable cancer cells (W-CIN+ cells) is the appearance of lagging chromosomes during anaphase, which is the result of erroneous and hyper-stable microtubule-kinetochore attachments ^9–11^. More recently, it was revealed that an abnormal increase in microtubule growth rates within mitotic spindles can be a direct trigger for the generation of lagging chromosomes and for W-CIN ^10,12–15^. In fact, increased microtubule growth seems to be a wide-spread mitotic defects present in W-CIN+ cancer cells ^10,13,15^. Significantly, restoration of this defect in various cancer cells was shown to be sufficient to suppress chromosome missegregation and W-CIN indicating a causality between increased microtubule polymerization and the induction of aneuploidy in cancer cells ^10,13,15^. Interestingly, in cancer cells aneuploidy is often accompanied with structural chromosome aberrations and *vice versa*, suggesting a link between W-CIN and S-CIN. Indeed, evidence for such a link was provided by demonstrating that W-CIN+ cells suffer from replication stress. Moreover, rescuing RS in these cancer cells resulted in suppression of W-CIN indicating that RS might link S-CIN to mitosis-mediated W-CIN ^16,17^. Mechanistically, it was demonstrated that moderate RS can cause premature centriole disengagement, which can contribute to spindle multipolarity in mitosis, thereby supporting missegregation of mitotic chromosomes ^18^. However, W-CIN+ cells exhibit only signs of very mild RS, which associates with increased mitotic microtubule growth rates leading to the generation of lagging chromosomes as a basis for W-CIN ^17^. Thus, there is clear evidence indicating that RS can affect mitotic chromosome segregation to cause W-CIN. However, the link between RS and mitotic defects is unknown.

RS can be caused by various means including DNA damage, abnormal DNA structures or shortage of replication factors or nucleotides ^4,6^. RS is prevalent in human cancer and pre-cancerous lesions and has been associated with S-CIN. In fact, oncogene activation including *MYC* or *CCNE1* amplification has been linked to the induction of RS and genome instability ^19–22^. Experimentally, inhibition of DNA polymerases using aphidicolin is widely used to induce RS, thereby allowing the induction of gradual levels of RS ^17^. Cells respond to severe RS by activating an intra-S phase checkpoint that involves the ATR kinase. ATR activation prevents the further progression of replication to allow DNA damage repair, but also stabilizes replication forks to allow subsequent re-start of replication ^23,24^. In contrast to severe RS that can lead to DNA damage and cell cycle arrest, W-CIN+ cancer cells were shown to exhibit only very mild RS, which can escape checkpoint control ^16,17^. These cells can further progress through the cell cycle and enter mitosis where under-replicated DNA might interfere with normal chromosome segregation ^25,26^.

For a normal DNA replication, human cells assemble ~500,000 pre-replication complexes (pre-RCs) in G1 phase by loading MCM helicase complexes (MCM2-7) and additional licensing factors onto specific chromatin sites, called origins of replication (ORCs). At the beginning of S phase, replication origin firing is triggered by CDC7 and CDK2 kinase activities that promote the recruitment of firing factors including GINS and CDC45 to form the active CDC45-MCM-GINS (CMG) helicase complex ^27–29^. During an unperturbed S phase, only ~10% of the licensed origins are fired indicating that the majority of licensed origins serves as back-ups. Indeed, upon RS, these dormant origins are activated leading to a higher origin density on chromatin (i.e. reduced inter-origin distances) ^30–33^. The mechanisms of dormant origin firing are not well understood, but several studies have revealed that S phase specific ATR inhibition is sufficient to induce dormant origin firing indicating that ATR limits origin firing during an unperturbed S phase ^34–37^. In this context, ATR acts as negative regulator of CDK1 during S phase, which negatively controls the assembly of the CDC7 counteracting the RIF1-PP1 protein phosphatase complex ^38–40^.

Importantly, upon RS or upon ATR-RIF1 inhibition in the absence of RS dormant origin firing is activated in a CDC7-dependent manner supporting the completion of DNA replication even when forks progress slowly ^27,30^. Thus, dormant origin firing seems to be beneficial for cells and is believed to suppress chromosomal instability during RS.

In contrast to this view, we found in this study that genes directly involved in replication origin firing are positively correlated with W-CIN in human tumor samples suggesting a role for increased origin firing in cancer chromosomal instability. We demonstrate that unscheduled induction of origin firing or dormant origin firing upon mild replication stress is sufficient to trigger W-CIN by increasing microtubule growth rates and chromosome missegregation in mitosis. Moreover, we show that chromosomally unstable cancer cells not only suffer from mild replication stress, but also exhibit increased origin firing leading to whole chromosome missegregation and W-CIN in these cancer cells.

## Results

### Genes involved in DNA replication origin firing are upregulated in human cancer and significantly correlate with W-CIN

To identify cancer-relevant genes that are associated with whole chromosomal instability (W-CIN) in human cancer we performed a systematic and comprehensive bioinformatic pan-cancer analysis using data from 32 different cancer types from *The Cancer Genome Atlas (TCGA)*. To quantify the degree of W-CIN in bulk tumor samples we used DNA copy number data and computed the whole genome integrity index (WGII) as a surrogate measure for W-CIN ^16,41^. To filter genes differentially expressed in W-CIN tumors, we divided the tumor samples into high and low WGII groups and compared their mean gene expression corrected for cancer type specific effects. Among the genes that positively correlate with the WGII score across most cancer types we found mitotic genes including *TPX2, RAE1, UBE2C, AURKA, AURKB, BUB1 and CDK1* (Fig. 1a). These candidates with functions in mitotic chromosome segregation are expected to be tightly associated with W-CIN and have indeed been identified previously as part of a CIN gene signature ^42^, thereby validating our systematic and unbiased bioinformatic approach. Our analysis also identified up-regulation of the known oncogenes *CCNE1* and *CCNE2* (encoding for cyclin E1/2) as being associated with W-CIN. *CCNE1* amplification has been previously linked to replication stress and genome instability ^19–22^. Interestingly, our analysis revealed an overall strong association of W-CIN with high expression of genes involved in DNA replication including *GINS1-4, CDC45, MCMs, DBF4, CDC7, RECQL4, PCNA, POLE* and *POLD2* (Fig. 1a). In fact, gene set enrichment analysis showed that genes positively associated with WGII scores are highly enriched for DNA replication factors (permutation test q-value = 0.00089, Fig. 1b). Moreover, a gene set annotated for DNA replication origin firing was found to be highly enriched at the top of all genes ranked by their correlation between WGII and expression (permutation test q-value = 8.04e−06, Fig. 1c) suggesting that high expression of genes involved in replication origin firing might be particularly associated with W-CIN. To investigate the association of origin firing gene expression including *GINS*, *MCM* and *CDC45* with W-CIN in individual cancer types we calculated Spearman correlation coefficients between expression and WGII scores. The strong correlation was reflected in many cancer types as shown in Fig. S1a.

**Figure 1:**
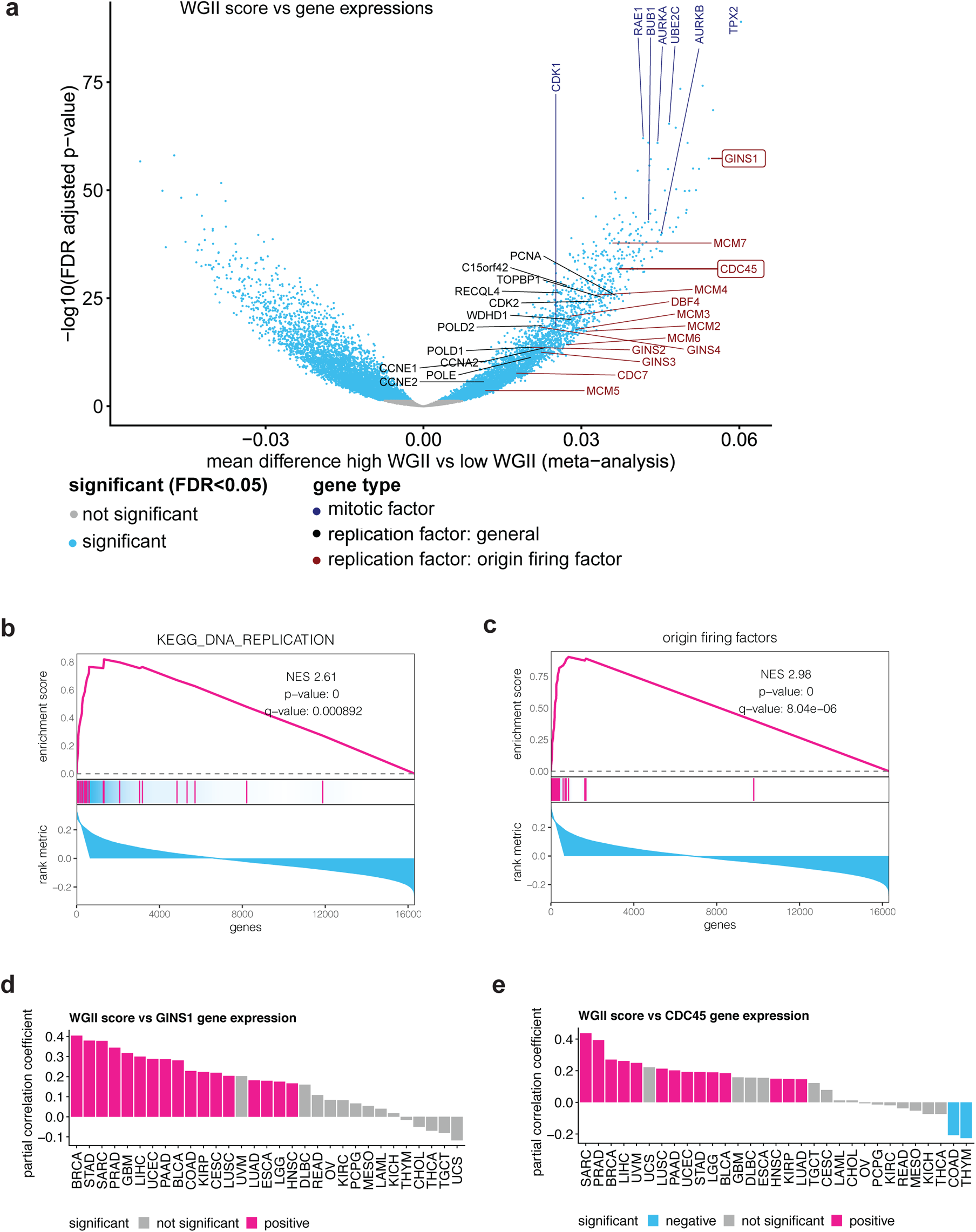
Positive association of genes involved in DNA replication origin firing with whole chromosome instability in human cancer specimens. **(a)** Association of gene expression and W-CIN in human cancer samples. The volcano plot shows the mean difference in normalized gene expression in tumor samples with high versus low WGII scores as a proxy measure for W-CIN. The WGII mean differences are adjusted for cancer type specific effects in the 32 different tumor types included in the pan-cancer analysis and the p-values are adjusted for multiple testing. **(b)** Gene set enrichment analysis for WGII-scores and genes involved in DNA replication. The analysis was performed using a gene set from KEGG annotated for DNA replication. The significance for the normalized enrichment score (NES) was evaluated by a permutation test and the pink bars indicate the position of DNA replication genes. **(c)** Gene set enrichment analysis for WGII-scores and genes associated with DNA replication origin firing. The analysis was performed using a set of manually curated origin firing factors as described in Material and Methods. The significance for the normalized enrichment score (NES) was assessed by a permutation test and the pink bars indicate the position of the origin firing genes. **(d)** *GINS1* gene expression is positively correlated with WGII scores in multiple cancer types, independent of the proliferation rate. The partial correlation coefficient between *GINS1* gene expression with WGII is shown, when the estimated proliferation rate is kept constant. **(e)** *CDC45* gene expression is positively correlated with WGII scores in multiple cancer types, independent of the proliferation rate as shown in (c).

Among the top genes whose expression correlate with W-CIN were *GINS1* and *CDC45*, both of which are well known key regulators of replication origin firing ^27^. Both, *GINS1* and *CDC45* expression showed a strong positive correlation with high WGII scores in various tumor entities, even when predicted proliferation rates ^43^ were taken into account suggesting that these origin firing genes might regulate W-CIN, but not overall proliferation in cancer specimens (Fig. 1d,e). Additionally, we found that copy number variations (CNVs) of many origin firing factors show an overall strong positive correlation with WGII scores and is most significant for *GINS1* (Fig. S1b). These results suggest that amplification of origin firing genes is a frequent event in various human cancers and correlates with high expression of these genes and W-CIN. Thus, based on our comprehensive pan-cancer analysis, we suggest that genes involved in origin firing represent potential oncogenes overexpressed in human cancer and might contribute to chromosomal instability.

### GINS1 or CDC45 overexpression increase replication origin firing without affecting replication fork progression

Our bioinformatic analysis identified the replication origin firing genes *GINS1* and *CDC45* as most significantly associated with W-CIN. To analyze the effects of high *GINS1* and *CDC45* expression on a cellular level and on genome stability, we stably overexpressed either *GINS1* or *CDC45* in chromosomally stable HCT116 cells that are characterized by proper chromosome segregation and DNA replication ^10,17^. We selected individual single cell clones for further analysis (Fig. 2a, Fig. S2a). First, we investigated how overexpression of the origin firing factors *GINS1* or *CDC45* affect DNA replication. For this, we performed DNA combing analysis upon DNA pulse labeling with nucleoside analogues 5-chloro-2ʹ-deoxyuridine (CldU) and 5-iodo-2ʹ-deoxyuridine (IdU) (Fig. 2b). Interestingly, *GINS1* or *CDC45* overexpression did not grossly affect the replication fork progression rate when compared to parental HCT116 cells (Fig. 2c, Fig. S2b). However, it significantly decreased the inter-origin distance demonstrating increased origin firing upon *GINS1* or *CDC45* overexpression (Fig. 2d, Fig. S2c). Origin firing at the beginning of S phase requires CDC7-mediated phosphorylation ^27,28,44^. Consequently, we found that inhibition of the CDC7 kinase using low concentrations of the small-molecule inhibitor XL-413 ^45^, which do not abrogate DNA replication, S phase progression or proliferation, fully restored proper inter-origin distances and thus, suppressed abnormally increased origin firing (Fig. 2d). Interestingly, CDC7 inhibition also slightly improved fork progression, which might be due to increased availability of nucleotides when normal levels of origin firing are restored in *GINS1* overexpressing cells (Fig. 2c). Together, *GINS1* or *CDC45* overexpression is common in human cancer and selectively increases replication origin firing without affecting DNA replication and fork progression *per se*.

**Figure 2:**
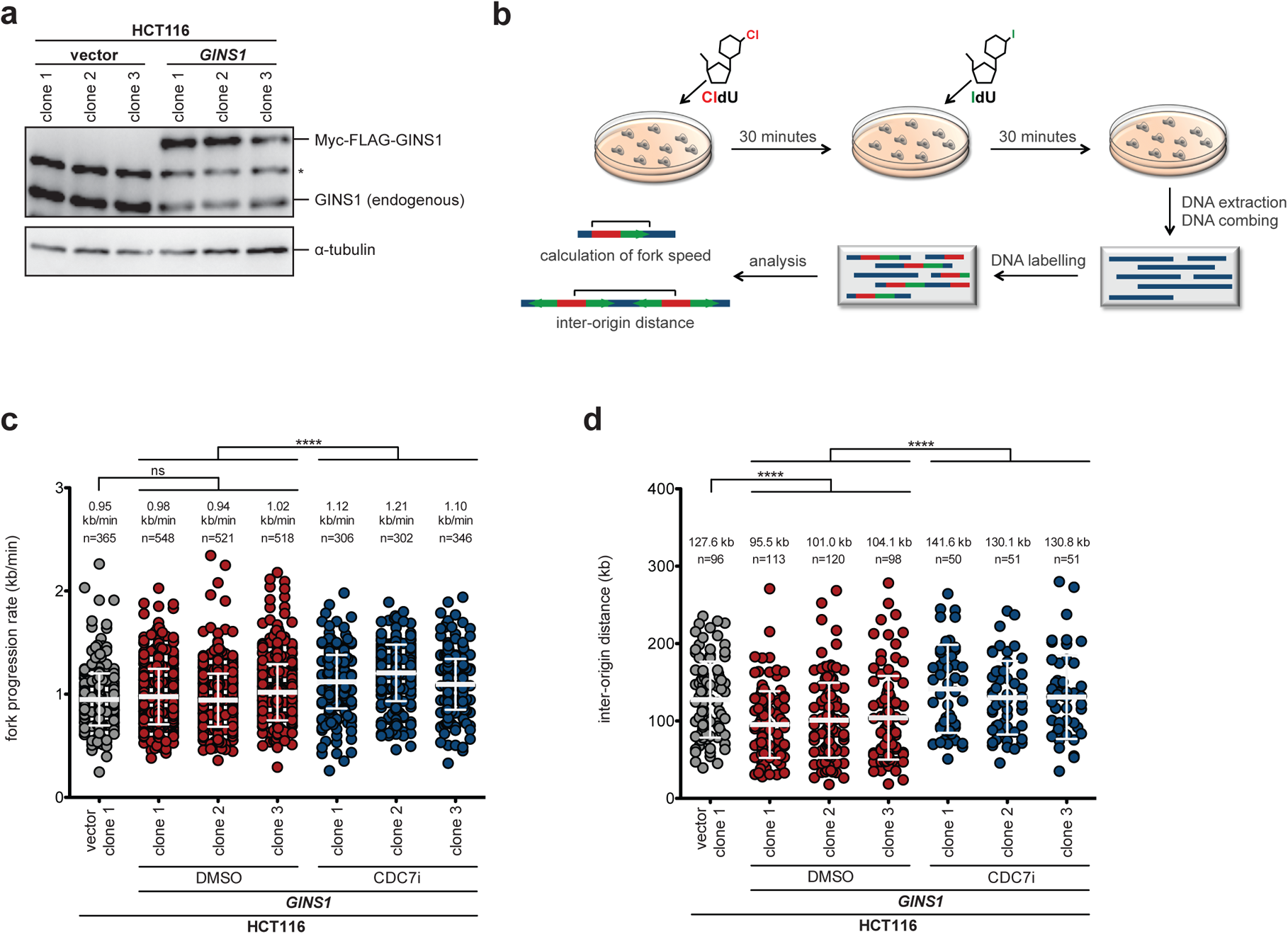
*GINS1* overexpression increase replication origin firing without affecting replication fork progression. **(a)** Generation of chromosomally stable HCT116 cells with stable GINS1 overexpression. A representative Western blot shows the expression of endogenous and overexpressed Myc-FLAG-tagged *GINS1* in three independent HCT116-derived single cell clones. Single cell clones transfected with empty vector serve as a control. α-tubulin was used as loading control. Star indicates an unspecific protein band. **(b)** Scheme illustrating DNA combing to determine replication fork progression and inter-origin distances as a measure for origin firing activity. Cells are pulse-labelled with 100 μM 5-chloro-2ʹ-deoxyuridine (CldU) and 100 µM 5-iodo-2ʹ-deoxyuridine (IdU) for 30 min each. DNA combing and subsequent detection of the newly synthesized DNA stretches allows the calculation of DNA replication fork speed and inter-origin distance. **(c)** Determination of replication fork progression rates in cells with or without *GINS1* overexpression and additional CDC7 inhibition. The indicated cell lines were pre-treated with 1 µM CDC7 inhibitor XL-413 (CDC7i) or DMSO as a control for 1 h before pulse-labelling with nucleoside analogues. Scatter dot plots show values for fork progression rates (mean ± SD, *t*-test). **(d)** Determination of origin firing frequency in cells with or without *GINS1* overexpression and additional CDC7 inhibition. Scatter dot plots show values for inter-origin distances (mean ± SD, *t*-test).

### Increased replication origin firing upon GINS1 or CDC45 expression causes W-CIN

Previous work showed that W-CIN+ cancer cells characterized by perpetual chromosome missegregation suffer from replication stress ^16,17^. Moreover, it has been demonstrated that chromosome missegregation and W-CIN in these cancer cells are triggered by abnormally increased microtubule growth rates during mitosis ^10,12,13,17^. Therefore, we evaluated whether increased origin firing triggers increased microtubule growth rates in mitosis leading to chromosome missegregation. Indeed, EB3-GFP tracking experiments in living mitotic cells revealed that overexpression of *GINS1* or *CDC45* was sufficient to cause increased mitotic microtubule growth rates (Fig. 3a; Fig. S2d) to a level typically detected in chromosomally instable cancer cells ^10,13,17^. Concomitantly, we detected a clear induction of lagging chromosomes during anaphase indicative for whole chromosome missegregation in cells with *GINS1* or *CDC45* overexpression (Fig. 3b, Fig. S2e). Importantly, chromosome missegregation was suppressed upon restoration of proper microtubule growth rates by low doses of Taxol (Fig. 3b; Fig. S2e), which was shown to correct abnormal microtubule growth rates in cancer cells ^10^. Moreover, microtubule growth rates and lagging chromosome were also suppressed upon CDC7 inhibition using XL-413 (Fig. 3a,b; Fig. S2d,e) demonstrating that chromosome missegregation is not only dependent on increased microtubule growth rates, but also on increased origin firing upon *GINS1* or *CDC45* overexpression. We therefore tested whether *GINS1* or *CDC45* expression is sufficient to induce W-CIN. For this, we analyzed single cell clones that were grown for 30 generations and determined the proportion of cells harboring chromosome numbers deviating from the modal number of 45 chromosomes (Figure 3c). These karyotype analysis indicate that overexpression of *GINS1* or *CDC45* is sufficient to cause the induction of aneuploidy and thus, of W-CIN (Fig. S2f, Fig. S3a,b).

**Figure 3:**
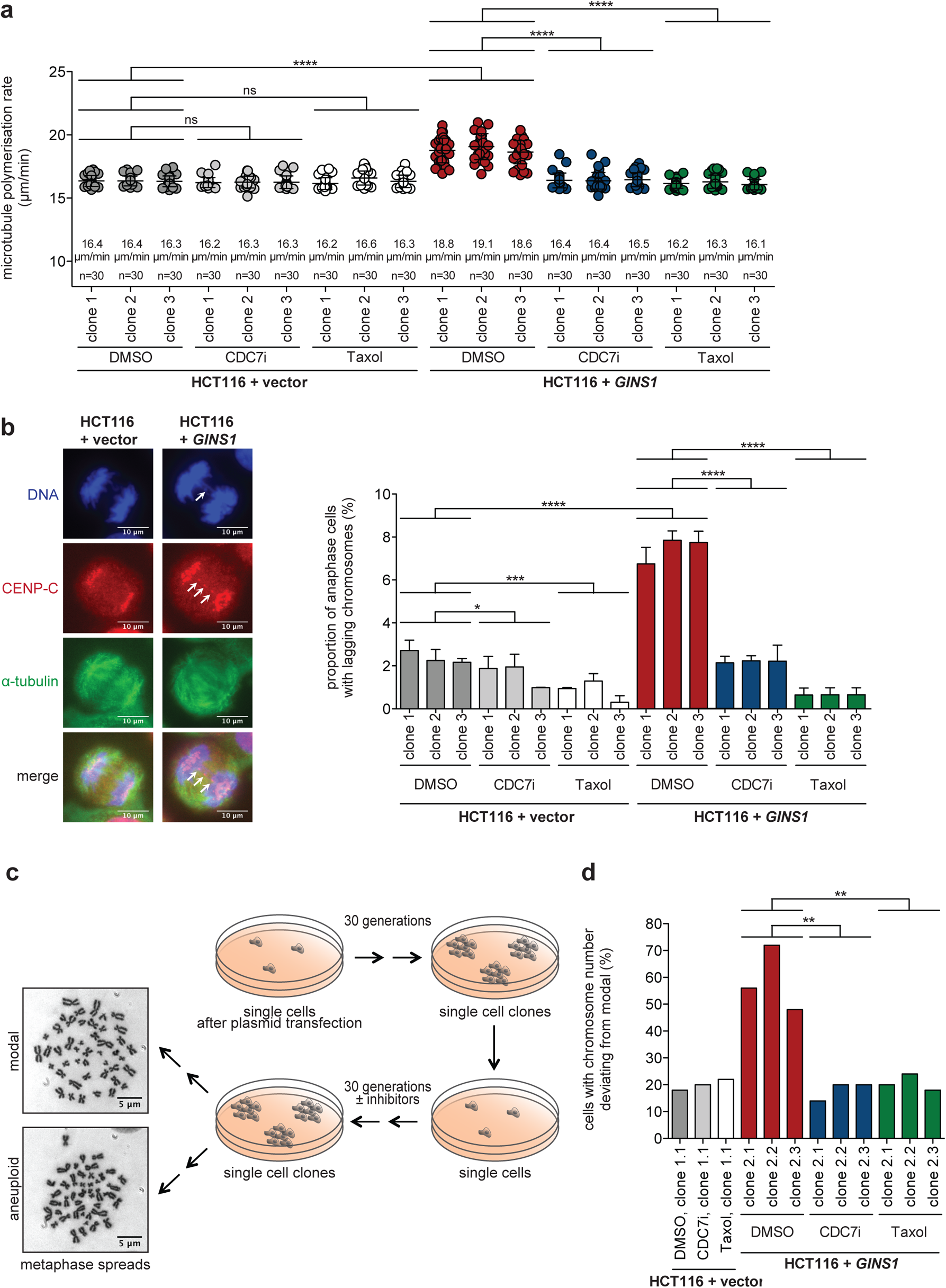
Overexpression of *GINS1* results in increased microtubule polymerization rates, chromosome missegregation and W-CIN. **(a)** Determination of mitotic microtubule growth rates in cells with or without overexpression of *GINS1* and in the presence or absence of CDC7 inhibition or Taxol treatment. The indicated single cell clones were treated with 1 µM of the CDC7 inhibitor XL-413 (CDC7i), or with 0.2 nM Taxol for 16 h and microtubule growth rates were determined in mitotic cells. Scatter dot plots show average microtubule growth rates (20 microtubules/cell, n=30 mitotic cells, mean ± SD, *t*-test). **(b)** Quantification of anaphase cells showing lagging chromosomes upon GINS1 overexpression. The indicated cell clones were treated as in (a) and the proportion of cells with lagging chromosomes was determined. Representative images of anaphase cells with or without lagging chromosomes (white arrows) are shown (scale bar: 10 µm). The bar graph shows the quantification of cells with lagging chromosomes (n≥300 anaphase cells from three to five independent experiments, mean ± SD, *t*-test). **(c)** Scheme illustrating the generation of single cell clones for karyotype analyses as a measure for W-CIN. Representative images of chromosome spreads with a normal and an aberrant karyotype are shown and chromosomes were counted from single cells (scale bar: 5 µm). **(d)** Determination of the proportion of *GINS1* overexpressing cells showing aneuploidy. The indicated single cell clones were grown for 30 generations in the presence of DMSO, CDC7i or Taxol. The chromosome numbers per cell were determined from metaphase spreads. The bar graph shows the proportion of cells with a karyotype deviating from the modal (45 chromosomes in HCT116 cells; n=50 metaphase spreads, *t*-test).

Moreover, we grew single cell clones with *GINS1* overexpression and additional long-term treatment with DMSO, (control), low-dose Taxol (to restore proper microtubule growth rates) or with XL-413 (to suppress additional origin firing) and determined the evolved karyotype variability (Fig. 3c, Fig. S4a). Both, Taxol and CDC7 inhibition fully suppressed the evolvement of aneuploidy indicating that W-CIN upon *GINS1* overexpression is dependent on both, increased microtubule growth rates and increased origin firing (Fig. 3d, Fig. S4b). It is of note that we were not able to cultivate single cell clones in the continuous presence of 1.0 µM XL-413 that was used in transient experiments before, which might be due to intracellular accumulation of the inhibitor. Instead, we used 0.5 µM XL-413 in these long-term experiments, which was sufficient to restore normal microtubule growth rates similar to 0.2 nM Taxol treatment (Figure S4c). Taken together, these results demonstrate that increased origin firing induced by *GINS1 or CDC45* overexpression is sufficient to trigger W-CIN by increasing mitotic microtubule growth rates, which is typically seen in W-CIN+ cancer cells ^10,13^.

### ATR-CDK1-RIF1-regulated dormant origin firing causes mitotic chromosome missegregation

Recent work showed that ATR signaling limits origin firing by counteracting CDK1 activity during S phase, thereby allowing balanced action of CDC7 and its counteracting RIF1-PP1 phosphatase complex ^38,40^. Consequently, ATR inhibition results in unleashed CDK1 activity that inactivates RIF1-PP1 and fosters increased origin firing mediated by the CDC7 kinase ^38^ (Fig. 4a). Based on these previous findings, we pharmacologically inhibited ATR kinase activity and verified the activation of dormant origin firing in an CDK1 and CDC7 dependent manner by performing DNA combing analysis (Fig. S5). Importantly, the ATRi-mediated dormant origin firing resulted in an increase in microtubule growth rates and chromosome missegregation in mitosis, both of which were suppressed upon concomitant inhibition of CDK1 or CDC7 indicating that ATRi-induced mitotic errors are mediated by CDK1/CDC7-triggered origin firing (Fig. 4b,c). Moreover, directly increasing CDK1 activity by stable expression of a constitutive active *CDK1* mutant (CDK1-AF) ^13^ was sufficient to increase microtubule growth rates and chromosome missegregation, again in a CDK1- and CDC7-activity-dependent manner (Fig. 4d,e). This further supports the notion that ATR inhibition acts through increased CDK1 activity to induce origin firing. Since increased CDK1 activity is expected to result in inhibition of the RIF1-PP1 phosphatase to induce CDC7-mediated origin firing (Fig. 4a), we depleted RIF1 by siRNAs (Fig. 4f) and evaluated the effects on mitosis. In fact, loss of RIF1 mimicked ATR inhibition or CDK1 activation and increased microtubule growth rates and chromosome missegregation in mitosis, again in a CDC7-, but not CDK1-dependent manner (Fig. 4g,h). Thus, abrogation of the ATR-RIF1 axis through CDK1 activation causes dormant origin firing leading to mitotic chromosome missegregation and W-CIN.

**Figure 4:**
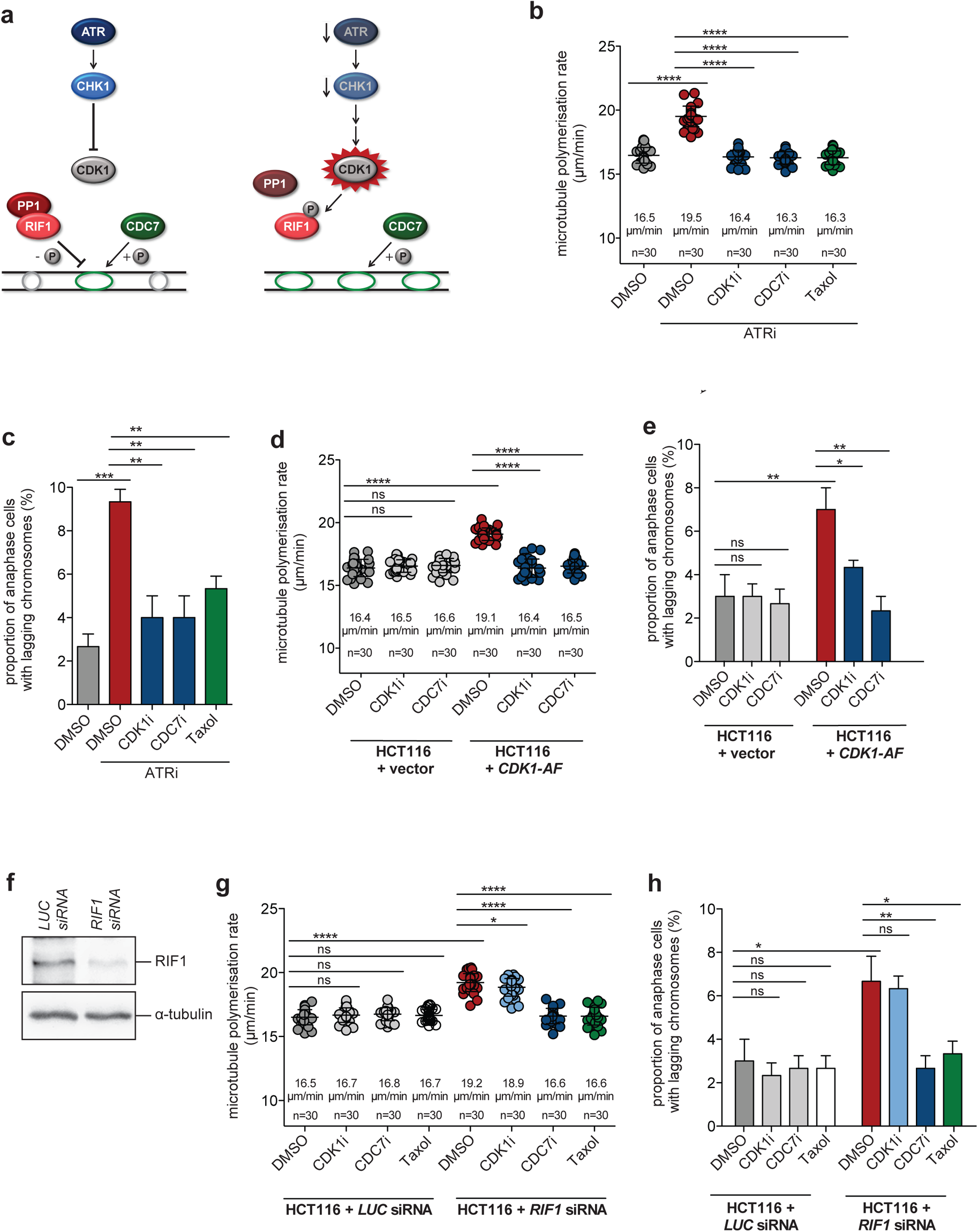
ATR-CDK1-regulated dormant origin firing causes mitotic chromosome missegregation. **(a)** Schematic illustrating the regulation of origin firing by ATR-CDK1 signaling. In unperturbed cells, ATR signaling limits CDK1 activity, which allows the balanced activity of the kinase CDC7 and the phosphatase complex RIF1-PP1. Upon ATR inhibition CDK1 activity increases and causes dissociation of the RIF1-PP1 complex resulting in CDC7-dependent origin firing (based on: Moiseeva *et al*., 2019c). **(b)** Determination of mitotic microtubule growth rates upon ATR inhibition-induced origin firing. HCT116 cells were treated with 1 µM ATR inhibitor ETP-46464 (ATRi) in combination with DMSO, 1 µM RO-3306 (CDK1i), 1 µM XL-413 (CDC7i), or 0.2 nM Taxol for 16 h. Scatter dot plots show average microtubule growth rates per cell (20 microtubules/cell, n=30 mitotic cells, mean ± SD, *t*-test). **(c)** Quantification of anaphase cells with lagging chromosomes after ATR inhibition-induced origin firing. Cells were treated as in (b) and the bar graph shows the proportion of anaphase cells with lagging chromosomes (n=300 anaphase cells, mean ± SD, *t*-test). **(d)** Measurements of mitotic microtubule growth rates in cells with or without expression of constitutive active CDK1. HCT116 cells stably expressing CDK1-AF were treated with CDK1i or CDC7i and scatter dot plots show average mitotic microtubule growth rates (20 microtubules/cell, n=30 mitotic cells, mean ± SD, *t*-test). **(e)** Quantification of anaphase cells with lagging chromosomes upon increased CDK1 activity and CDK1i or CDC7i treatment. Cells were treated as in (d) the incidence of lagging chromosomes in anaphase cells was determined (n=300 anaphase cells, mean ± SD, *t*-test). **(f)** siRNA-mediated downregulation of RIF1. HCT116 cells were transfected with siRNAs targeting *LUCIFERASE (LUC)* or *RIF1*. After 48 h, western blotting confirmed knockdown efficiency. α-tubulin levels were detected as loading control. **(g)** Measurements of mitotic microtubule growth rates in cells with or without downregulation of RIF1 and treatment with CDK1i, CDC7i or Taxol. After siRNA transfection cells were treated with CDK1i, CDC7i or Taxol for 16 h and microtubule growth rates were measured. Scatter dot-plots show average microtubule growth rates per cell (20 microtubules/cell, n=30 mitotic cells, mean ± SD, *t*-test). **(h)** Quantification of anaphase cells with lagging chromosomes after downregulation of RIF1 and treatment with CDK1i, CDC7i or Taxol. Cells were treated as in (g) and bar graphs show the proportion of anaphase cells with lagging chromosomes (n=300 anaphase cells, mean ± SD, *t*-test).

### Replication stress-induced dormant origin firing causes mitotic chromosome missegregation

W-CIN+ cancer cells suffer from mild replication stress that can be mimicked by treatment with very low concentrations (100 nM) of the DNA polymerase inhibitor aphidicolin ^16,17^. Replication stress is known to activate dormant origin firing as a compensation mechanism to complete DNA replication when replication forks progress too slowly ^46^. We asked whether dormant origin firing induced by cancer-relevant mild replication stress can cause whole chromosome missegregation in mitosis. To this end, we treated chromosomally stable HCT116 cells with 100 nM aphidicolin to induce mild replication stress and performed DNA combing analysis. As expected, aphidicolin reduced replication fork progression (Fig. 5a) and decreased the inter-origin distances indicating that dormant origin firing represents a consequence of slowed fork progression upon RS (Fig. 5b). Importantly, CDC7 or CDK1 inhibition did not affect the slowed fork progression rates, but fully restored normal inter-origin distances (Fig. 5a,b) indicating that partial CDK1 or CDC7 inhibition can selectively used to suppress dormant origin firing during aphidicolin-induced RS. Then we tested whether replication stress-induced dormant origin firing can trigger mitotic errors. As shown before ^17^, mild replication stress increased mitotic microtubule growth rates and lagging chromosomes (Fig. 5c,d). Importantly, these effects were fully suppressed when dormant origin firing was selectively inhibited upon CDK1 or CDC7 inhibition (Fig. 5c,d), which demonstrates that dormant origin firing during mild replication stress in S phase represents a trigger for whole chromosome missegregation during the subsequent mitosis.

**Figure 5:**
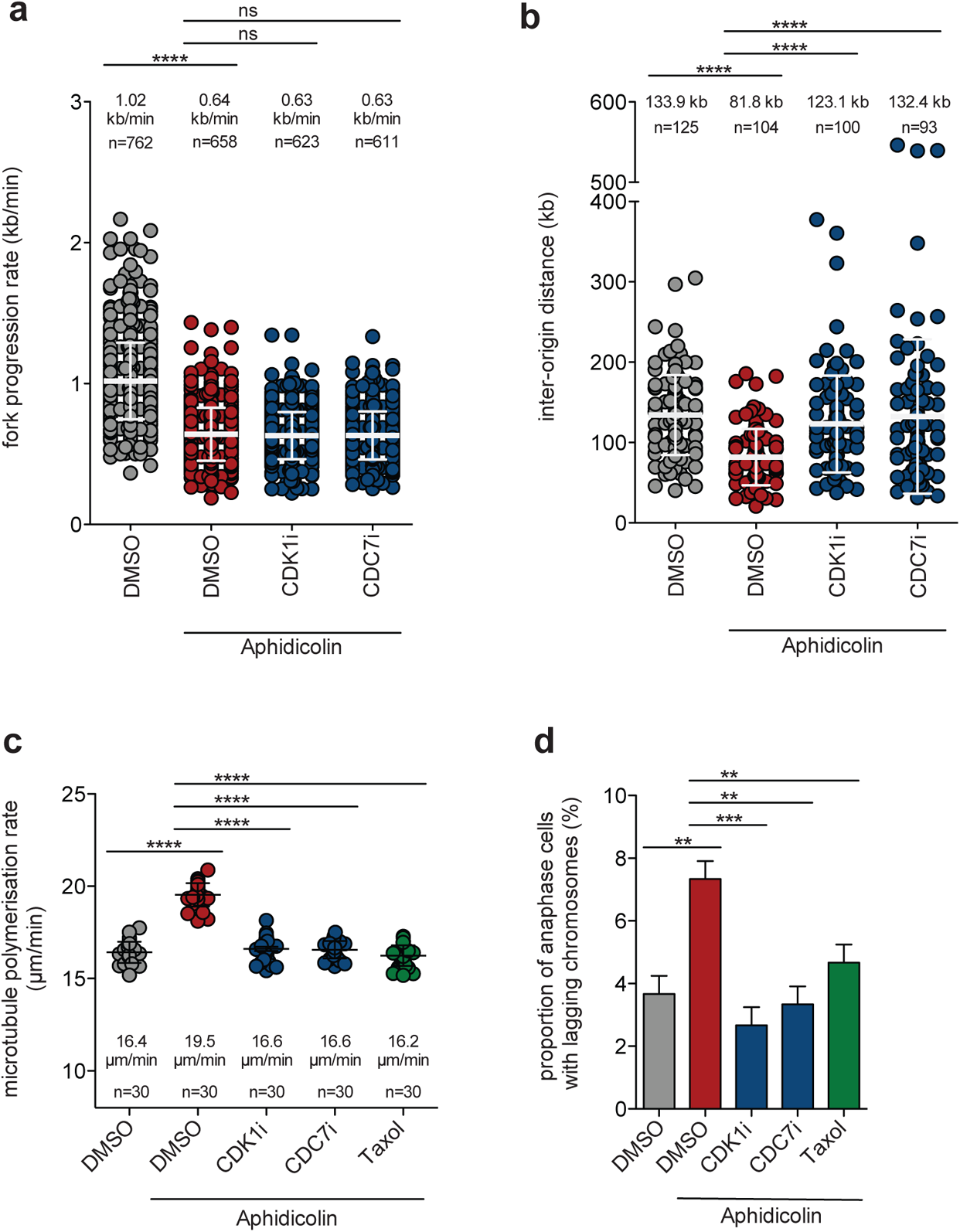
Replication stress-induced dormant origin firing causes mitotic chromosome missegregation. **(a)** Measurements of replication fork progression rates in chromosomally stable HCT116 cells upon mild replication stress and treatment with CDK1 or CDC7 inhibitors. Cells were treated with 100 nM aphidicolin to induce mild replication stress and additionally with DMSO, 1 µM RO-3306 (CDK1i) or 1 µM XL-413 (CDC7i) for 1 h. Subsequently, cells were subjected to DNA combing analysis and replication fork progression rates were determined (mean ± SD, *t*-test). **(b)** Measurements of inter-origin distances as a measure for origin firing frequencies. Cells were treated as in (a) and inter-origin distances were determined (mean ± SD, *t*-test). **(c)** Determination of mitotic microtubule growth rates upon mild replication stress and treatment with CDK1i or CDC7i. HCT116 cells were treated with 100 nM aphidicolin and CDK1i, CDC7i, or 0.2 nM Taxol for 16 hrs and microtubule growth rates were measured in living mitotic cells. Scatter dot plots show average microtubule growth rates per cell (20 microtubules/cell, n=30 mitotic cells, mean ± SD, *t*-test). **(d)** Quantification of anaphase cells showing lagging chromosomes after induction of mild replication stress and treatment with CDK1i, CDC7i or Taxol. Cells were treated as in (c) and the bar graph shows the proportion of cells with lagging chromosomes (n=300 anaphase cells, mean ± SD, *t*-test).

### Activation of dormant origin firing during early S phase triggers mitotic errors

To further investigate whether the ATR-CDK1-CDC7-dependent regulation of dormant origin firing acts during S phase to cause mitotic dysfunction we established a schedule for inhibitor treatments during different phases of the cell cycle prior to the analysis of mitotic phenotypes (Fig. 6a). We treated cells with ATRi only during a two-hour time window during early S phase followed by washout of the drug. This S phase-specific treatment was sufficient to increase microtubule growth rates and to induce lagging chromosomes in the subsequent mitosis (Fig. 6b,c). Moreover, the mitotic errors were only suppressed by CDK1 or CDC7 inhibition when applied also during early S phase, but not when applied at the G2/M transition (Fig. 6b,c) indicating that the ATRi-mediated increase in CDK1 and CDC7-mediated origin firing is required during early S phase to induce errors in the subsequent mitosis. This finding was further supported by using HCT116 cells with increased CDK1 activity (expressing CDK1-AF) where inhibition of CDK1 or CDC7 only during early S phase, but not in late S phase, G2 or at G2/M rescued the mitotic defects (Fig. 6d,e). Finally, we increased CDK1 activity in a cell cycle stage dependent manner by inhibiting the WEE1 kinase, a negative regulator of CDK1 ^47^. WEE1 inhibition was previously shown to induce dormant origin firing in a CDK1-dependent manner ^39,48^.

**Figure 6:**
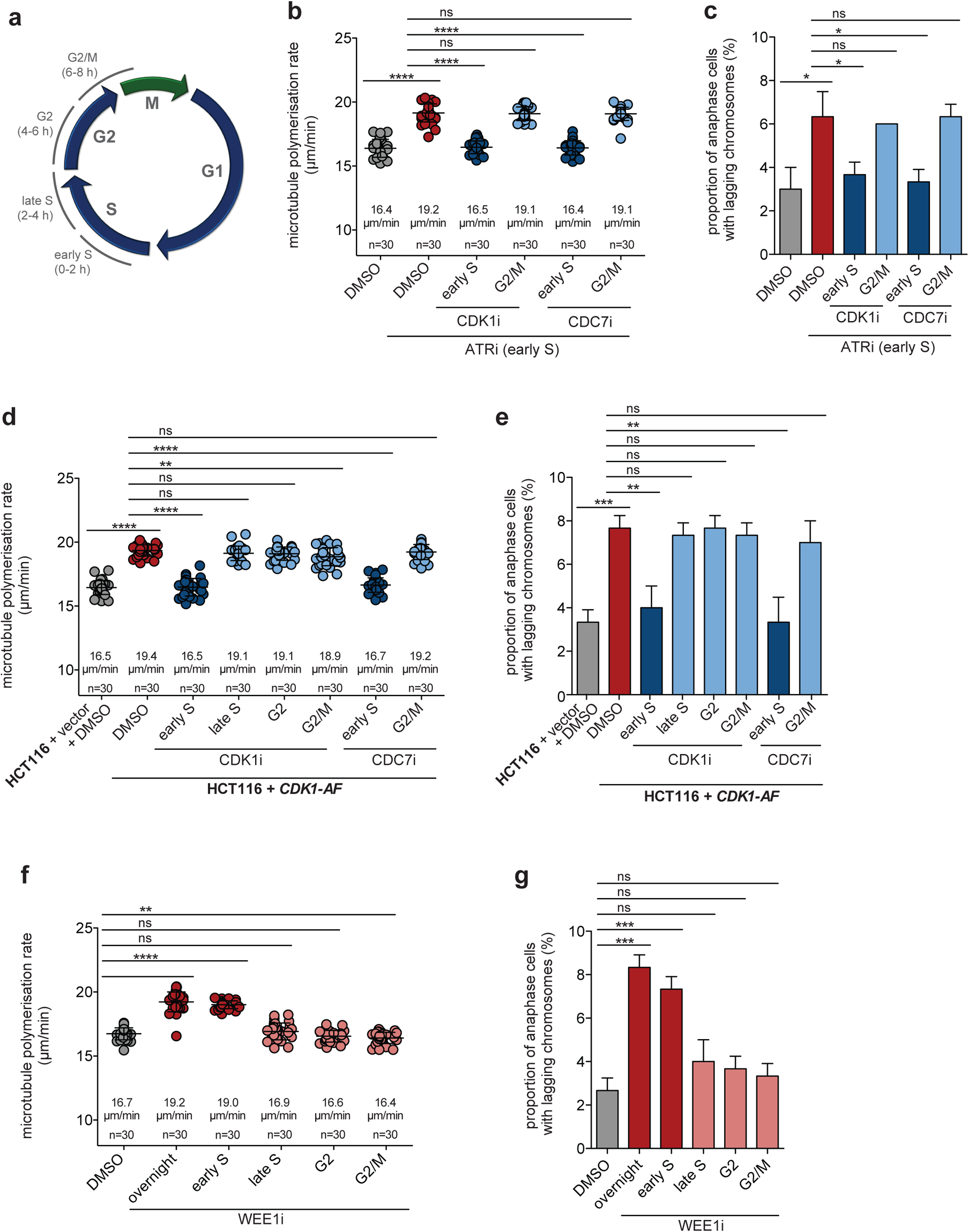
Activation of dormant origin firing specifically during early S phase triggers mitotic errors. **(a)** Depiction of cell cycle dependent treatment windows. Cells were treated at specific time points during the cell cycle and the effects were evaluated during the subsequent mitosis. **(b)** Measurements of mitotic microtubule growth rates in HCT116 cells with S phase-specific ATR inhibition (1.0 µM ETP-46464, ATRi) and additional CDK1 inhibitor (1.0 µM RO-3306, CDK1i) or CDC7 inhibitor (1.0 µM XL413, CDC7i) treatment during the indicated time windows. All drugs were washed-out after 2 h treatment and microtubule growth rates were measured in mitosis. Scatter dot-plots show average microtubule growth rates per cell (20 microtubules/cell, n=30 mitotic cells, mean ± SD, *t*-test). **(c)** Quantification of anaphase cells with lagging chromosomes after cell cycle specific drug treatments as used in (b). The proportion of anaphase cells with lagging chromosomes was determined (n=300 anaphase cells, mean ± SD, *t*-test). **(d)** Measurements of mitotic microtubule growth rates in cells with elevated CDK1 activity (CDK1-AF) and treatment with CDK1i or CDC7i during the indicated time windows. Scatter dot plots show average microtubule growth rates per cell (20 microtubules/cell, n=30 mitotic cells, mean ± SD, *t*-test). **(e)** Quantification of anaphase cells with lagging chromosomes using CDK1-AF expressing cells with or without cell cycle specific CDK1i and CDC7i treatment as used in (d). The proportion of anaphase cells with lagging chromosomes was determined (n=300 anaphase cells, mean ± SD, *t*-test). **(f)** Measurements of mitotic microtubule growth rates in cells treated with 75 nM of the WEE1 inhibitor MK-1775 (WEE1i) for 2 hrs during the indicated cell cycle phases. Scatter dot-plots show average microtubule growth rates per cell (20 microtubules/cell, n=30 mitotic cells, mean ± SD, *t*-test). **(g)** Quantification of anaphase cells with lagging chromosomes after cell cycle specific WEE1i treatment as used in (f). The proportion of anaphase cells with lagging chromosomes was determined (n=300 anaphase cells, mean ± SD, *t*-test).

Significantly, WEE1 inhibition led to an increase in mitotic microtubule growth rates and to an induction of lagging chromosomes only when applied during a two-hour time window in early S phase, but not in late S phase, G2 or at G2/M (Fig. 6f,g). Thus, dormant origin firing, specifically during early S phase and either triggered upon mild replication stress or upon ATR inhibition or CDK1 activation, is sufficient to cause mitotic defects leading to whole chromosome missegregation and W-CIN.

### Dormant origin firing is a trigger for W-CIN in chromosomally unstable cancer cells

Chromosomally unstable, aneuploid colorectal cancer cells (W-CIN+ cells) are characterized by increased mitotic microtubule growth rates, increased incidence of lagging chromosomes and by mild replication stress ^10,13,16,17^. We asked whether dormant origin firing represents a trigger for W-CIN in these cancer cells. To this end, we performed DNA combing analysis using three different W-CIN+ cell lines in the presence or absence of CDC7 inhibition. In line with previous work ^16,17^, we found that the W-CIN+ cells showed decreased replication fork progression when compared to chromosomally stable HCT116 cells, which was largely unaffected by CDC7 inhibition (Fig. 7a). Moreover, all W-CIN+ cell lines showed increased dormant origin firing reflected by decreased inter-origin distances that was suppressed upon CDC7 inhibition (Fig. 7b), indicating that CDC7 inhibition can be used to discriminate between slow fork progression and increased origin firing in W-CIN+ cancer cells. As shown before ^10,13,17^, W-CIN+ cancer cells exhibit increased mitotic microtubule growth rates that cause the generation of lagging chromosomes (Fig. 7c,d). Importantly, both, abnormal microtubule growth rates and the generation of lagging chromosomes were suppressed upon restoration of proper origin firing after CDC7 inhibition (Fig. 7c,d) indicating that increased origin firing, but not slowed replication fork progression acts as a trigger for subsequent mitotic errors. It is of note that we recently showed that perpetual chromosome missegregation in W-CIN+ cells is suppressed upon CDK1 inhibition ^13^, which is in line with our results presented here showing that CDK1 unleashed upon ATR inhibition increased origin firing (Fig. 5). To further support our findings, we partially depleted either CDC7 or different components of the CMG helicase (GINS1, CDC45 and MCM2), all of which are well-known to influence dormant origin firing ^33,49^, in W-CIN+ cells (Fig. S6) and analyzed microtubule growth rates and chromosome segregation in mitosis. Similar to CDC7 or CDK1 inhibition, siRNA-mediated partial knockdown of *CDC7*, *GINS1*, *CDC45* or *MCM2* restored normal mitotic microtubule polymerization rates and chromosome segregation in all three W-CIN+ cell lines (Fig. 7e,f). Thus, dormant origin firing in chromosomally unstable cancer cells suffering from mild replication stress acts as a trigger for subsequent mitotic chromosome missegregation and chromosomal instability.

**Figure 7:**
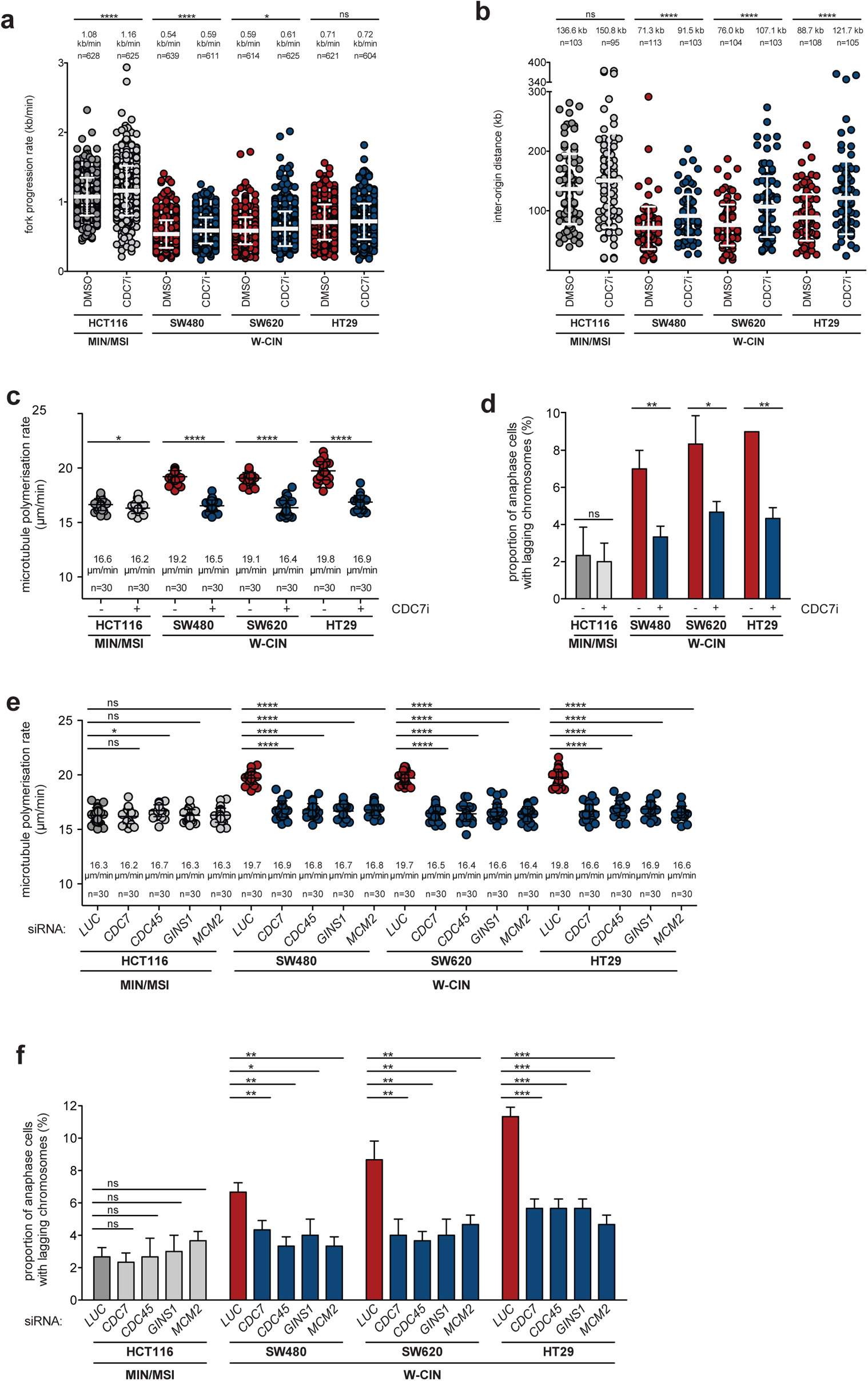
Dormant origin firing is a trigger for W-CIN in colorectal cancer cells. **(a)** Measurements of replication fork progression rates in different W-CIN+ colorectal cancer cell lines in the presence or absence of CDC7i. The indicated cell lines were treated with CDC7i for 2 h and subjected to DNA combing analysis and replication fork progression rates were determined (mean ± SD, *t*-test). **(b)** Measurements of inter-origin distances as a measure for origin firing frequencies. The different cell lines were treated as in (a) and inter-origin distances were determined (mean ± SD, *t*-test). **(c)** Measurements of mitotic microtubule growth rates in different CIN+ cells treated with CDC7i. The indicated colorectal cancer cell lines were treated with CDC7i for 16 h and microtubule growth rates were determined in mitotic cells. Scatter dot plots show average microtubule growth rates per cell (20 microtubules/cell, n=30 mitotic cells, mean ± SD, *t*-test). **(d)** Proportion of W-CIN+ cells with lagging chromosomes after CDC7i treatment. The indicated cell lines were treated as in (c) and the proportion of anaphase cells with lagging chromosomes was determined (n=300 anaphase cells, mean ± SD, *t*-test). **(e)** Measurements of mitotic microtubule growth rates in W-CIN+ cells after downregulation of CDC7 or CMG components. The indicated cancer cell lines were transfected with siRNAs targeting *CDC7*, *CDC45*, *GINS1*, or *MCM2*. LUCIFERASE (LUC) siRNA was sued a control. 48 hrs after transfection microtubule growth rates were determined in mitotic cells. Scatter dot plots show average microtubule growth rates per cell (20 microtubules/cell, n=30 mitotic cells, mean ± SD, *t*-test). **(f)** Proportion of W-CIN+ cells with lagging chromosomes after downregulation of CMG components. The indicated cell lines were transfected as in (e) and the proportion of anaphase cells with lagging chromosomes was determined (n=300 anaphase cells, mean ± SD, *t*-test).

## Discussion

This study revealed that increased replication origin firing can act as a so far unrecognized trigger for mitotic chromosome missegregation and the induction of whole chromosome instability (W-CIN) in human cancer cells. Origin firing-induced W-CIN involves an induction of abnormally increased microtubule growth rates in mitosis, which is known to cause W-CIN ^10,15,17^. Induction of origin firing occurs in different scenarios: (i) upon overexpression of potentially oncogenic origin firing genes causing dormant origin firing associated with W-CIN in human cancer specimens, (ii) experimentally, by using the DNA polymerase inhibitor aphidicolin, which is known to induce mild replication stress and dormant origin firing, (iii) upon inhibition of the ATR-RIF1 axis known to negatively regulate dormant origin firing during an unperturbed S phase ^30,38^, and (iv) in W-CIN+ cancer cells known to exhibit endogenous mild replication stress ^16,17^. In all cases, we found that increased origin firing, but not replication stress *per se*, is sufficient to induce mitotic chromosome missegregation and W-CIN.

Origin firing requires the licensing of origins in G1 phase and is initiated at the beginning of S phase by CDK- and CDC7-mediated phosphorylation and assembly of the CDC45-MCM-GINS (CMG) helicase complex ^49^. In human cells, there is a large excess of licensed over fired origins. During an unperturbed DNA replication most origins remain dormant, but during replication stress dormant origins can fire and this is thought to represent a compensatory mechanism to rescue RS ^30,46^. Our DNA combing results support this view and showed that even mild RS, which is not sufficient to activate the ATR-dependent checkpoint ^17^, induces dormant origin firing. Importantly, CIN+ cancer cells not only show slowed replication fork progression, but are also characterized by dormant origin firing. The causal link between replication stress and increased dormant origin firing is well established. In fact, partial depletion of MCM2-7 complexes, which only impairs dormant origin firing during replication stress, but not normal DNA replication timing, results in an induction of markers for under-replicated DNA including DNA damage, mitotic DNA synthesis, micronuclei formation and formation of 53BP1 nuclear bodies ^31,33^. Therefore, it was concluded that dormant origin firing is beneficial for cells, rescues replication stress and possibly, suppresses chromosomal instability ^30^. However, our data presented here clearly indicate that dormant origin firing can contribute to chromosomal instability by triggering mitotic errors.

It is currently not well understood how dormant origin firing is initiated during replication stress. Possibly, licensed dormant origins are passively removed during unperturbed DNA replication. Consequently, a subset of dormant origins might not be removed during RS due to the slowly progressing forks and are allowed to fire ^30,46^.

On the other hand, it has been demonstrated that inhibition of ATR, resulting in activation of CDK1 and abrogation of the RIF1-PP1 phosphatase complex in S phase, is sufficient to induce dormant origin firing in the absence of replication stress ^34,37–39^. This suggests that a non-checkpoint pool of ATR that is active during an unperturbed S phase can limit origin firing. Our results support this model and showed that ATR inhibition, CDK1 activation or loss of RIF1 results in increased origin firing and leads to subsequent mitotic dysfunction and chromosome missegregation in an origin firing dependent manner.

The intriguing link between increased origin firing and increased mitotic microtubule growth rats, which is responsible for chromosome missegregation in mitosis, is currently not understood. One can speculate that unscheduled origin firing might activate yet unknown signaling pathways leading to deregulation of microtubule associated proteins. In fact, the processive microtubule polymerase ch-TOG might be a relevant target since it has been demonstrated that its overexpression, observed in various cancers ^50,51^, is sufficient to increase microtubule growth rates and to induce whole chromosome missegregation ^10,52^. In addition, other microtubule plus end binding proteins with functions in microtubule plus-tip assembly ^53^ might also be subject to functional modulation in response to increased origin firing in S phase.

Comprehensive proteomic approaches could provide important clues on microtubule associated proteins that might be deregulated specifically after increased origin firing. Intriguingly, our cell cycle dependent analysis revealed that modulation of origin firing specifically during early S phase, but not in late S phase or G2 is required to mediate the subsequent mitotic errors and W-CIN. Thus, there is a time window of origin firing during early S phase, which is of particular importance for chromosomal instability. It is well known that DNA replication has a complex and distinct spatio-temporal organization ^54^. Late replicating domains often show low origin densities, which might contribute to their under-replication in response to replication stress. In fact, these regions were identified as common fragile sites (CFSs), which are prone to fragility and represent common breakpoints in cancer cells ^54,55^. In contrast, the recently discovered early-replicating fragile sites (ERFS) are located in early replicating chromosome domains and contain highly expressed genes and a higher origin density ^56^. These early replicating chromosome domains seem to be highly cancer relevant. More than 50% of all translocations in B-cell lymphomas were found to be associated with ERFSs ^56^. Our results now indicate that mitotic errors are more likely to result from increased origin firing in early S phase, i.e. in early replicating domains. Whether this is directly linked to ERFSs or whether transcription-replication conflicts, which might be more prevalent upon increased origin firing in early replicating domains ^57^ remains to be shown. Overall, these new results might suggest mechanistic links between S-CIN affecting early replicating chromosome domains and W-CIN affecting whole chromosomes.

It is well known that cancer cells, in particular W-CIN+ cells, suffer from mild replication stress, which can be caused by various means including DNA damage, nucleotide or replication factor shortage and oncogene expression ^4,6,16,17,58^. The latter might be of particular relevance in cancer. For instance, overexpression of *CCNE1* (encoding for cyclin E) or *MYC* has not only been linked to RS, but, interestingly, also to increased origin firing ^58^. The additionally fired origins were associated with collapse of replication forks leading to DNA damage, thereby linking oncogene-induced origin firing to chromosomal rearrangements and thus, to S-CIN ^22^. Intriguingly, both, ERFSs and oncogene-induced origins map to highly transcribed chromosomal domains suggesting a possible role of transcription-replication conflicts in CIN ^57^. Interestingly, previous studies also demonstrated that high expression of oncogenes like *CCNE1* or *MYC* can also interfere with proper chromosome segregation in mitosis, but the underlying mechanisms remained unclear ^19,21,59,60^. Based on our work presented here, it seems plausible that oncogenes affect mitosis and induce W-CIN through their role in inducing origin firing.

In addition to the classical oncogenes, our systematic pan-cancer analysis identified origin firing genes itself as putative oncogenes that increase origin firing and induce mitotic errors. We found that *GINS1*, *CDC45*, *MCMs* and others are frequently upregulated in various human cancer types and their high expression correlate significantly with W-CIN. Similar to mitotic genes that are known to influence mitotic chromosome segregation directly (e.g. *AURKA*, *TPX2* located on chromosome 20q; ^42,61^) we found that high expression of origin firing genes like *GINS1* were associated with copy number gains across many different cancer types indicating that amplification of origin firing genes is frequent in human cancer. Importantly, we showed that overexpression of *GINS1* or *CDC45* alone is sufficient to trigger dormant origin firing without inducing replication stress *per se*, i.e. without altering replication fork velocity. This specific induction of origin firing was nevertheless sufficient to cause mitotic chromosome missegregation, aneuploidy and W-CIN demonstrating that origin firing, but not slowed replication kinetics is responsible for mitotic dysfunction and W-CIN. Since W-CIN has been linked to tumor progression, tumor aggressiveness and therapy resistance ^1,2^, it is not surprising that high expression of *GINS1* or *CDC45* was found to be associated with poor prognosis in different tumor types supporting putative oncogenic functions of genes involved in origin firing ^62,63^.

## Material and methods

### Cell culture

HCT116, HT29, SW480, and SW620 cells were obtained from ATCC (USA). Cells were cultivated in RPMI1640 medium (PAN-Biotech GmbH, Germany) supplemented with 10 % fetal bovine serum (FBS; Corning Inc., USA), 100 units/ml penicillin, and 100 μg/ml streptomycin (Anprotec, Germany). HCT116 + *CDK1-AF* and the corresponding control cells^13^ were grown in medium with 300 µg/ml G418 (Santa Cruz, USA). All cells were grown in a humidified atmosphere at 37 °C and 5 % CO_2_.

### Plasmid and siRNA transfections

For EB3-GFP tracking experiments, cells were transfected with 10 µg pEGFP-*EB3* (kindly provided by Linda Wordeman, Seattle, WA, USA) using a GenePulser Xcell (Bio-Rad Laboratories, USA) at 500 µF and 300 V (HCT116, SW620), or 950 µF and 220 V (SW480, HT29). Cells were transfected with siRNAs (60 pmol; Sigma-Aldrich, Germany) using ScreenFect®siRNA (ScreenFect GmbH, Germany) or Lipofectamine RNAiMAX (Thermo Fisher Scientific, USA) according to the manufacturer protocols. The used siRNA sequences are listed below. Further experiments were performed 48 hrs after transfection and Western blotting was used to confirm transfection efficiency.

*LUCIFERASE (LUC)*: 5’-CUUACGCUGAGUACUUCGAUU-3’;

*CDC45*: 5’-UUCAUCCAGGCUCUGGACAGC-3’; *CDC7*: 5’-AAGCUCAGCAGGAAAGGUG-3’; *GINS1*: 5’-AAAGAUCUCUUGCUACUUAdTdT-3’;

*MCM2*: 5’GGAGCUCAUUGGAGAUGGCAUGGAA-3’;

*RIF1*: 5’-AAGAGCAUCUCAGGGUUUGCUdTdT-3’

### Generation of stable cell lines

For the generation of HCT116-derived cell lines stably expressing *CDC45* or *GINS1*, HCT116 cells were transfected with 0.75 µg or 1.5 µg mCherry-*CDC45* (kindly provided by Helmut Pospiech, FLI, Jena, Germany ^64^) and 1.5 µg or 2.0 µg pCMV6-Myc-FLAG-*GINS1* (OriGene Technologies, Inc., USA), respectively, using METAFECTENE (Biontex, Germany) according to the manufacturer instructions. Several single cell clones were grown in medium supplemented with 300 µg/ml G418 (Santa Cruz, USA) and selected for further analysis.

### Cell treatments

To restore proper microtubule polymerization rates, cells were grown in the presence of 0.2 nM Taxol (Sigma-Aldrich, Germany) as shown before^10,12^. The inhibitors ETP-46464 (1.0 µM; Selleck Chemicals, USA), MK-1775 (75 nM; Selleck Chemicals, USA), RO-3306 (1.0 µM; Santa Cruz, USA), and XL-413 (0.5-1.0 µM; Tocris Bioscience, UK) were used to inhibit ATR, WEE1, CDK1, and CDC7 kinases, respectively. All inhibitors were titrated to ensure that cell cycle progression was not affected. Cells were treated with 100 nM aphidicolin (Santa Cruz, USA) to induce mild replication stress as described before ^17^. Corresponding volumes of DMSO or H_2_O were used as controls.

### Analysis of microtubule polymerization rates

EB3-GFP tracking experiments were performed to determine microtubule polymerization rates^10,65^. 48 hrs after transfection with pEGFP-*EB3*, cells were treated with 2.0 µM Dimethylenastron (DME; Calbiochem, USA) for 1-2 hrs to accumulate cells in prometaphase^10^. To visualize microtubule plus tips, live cell microscopy was performed using a DeltaVision Elite microscope (GE Healthcare, UK) equipped with a PCO Edge sCMOS camera (PCO, Germany) and the softWoRx® 6.0 Software Suite (GE Healthcare, USA). Mitotic cells were monitored for 30 seconds in total, and images were taken every 2 seconds. During image acquisition, cells were incubated at 37 °C and 5 % CO_2_. The softWoRx® 6.0 Software Suite (GE Healthcare, USA) was used for image deconvolution and analysis. Average microtubule growth rates were calculated from 20 microtubules per cell.

### Quantification of anaphase cells exhibiting lagging chromosomes

Cells were synchronized in anaphase by a double thymidine block followed by a release for 8.5-9.5 hrs ^13^. Cells were fixed with 2 % paraformaldehyde/PBS for 5 minutes and then with ice-cold 100 % methanol for 5 minutes at −20 °C. To visualize microtubules, kinetochores, and the DNA, cells were stained with anti-α-tubulin (1:700, B-5-1-2, Santa Cruz, USA, cat no sc-23948), anti-CENP-C (1:1000, MBL International Corporation, USA, cat no PD030) and secondary antibodies conjugated to Alexa-Fluor488 (1:1000, Thermo Fisher Scientific, USA, cat no A-11029) and Alexa-Fluor594 (1:1000, Thermo Fisher Scientific, USA, cat no A-11076), and Hoechst33342 (1:15000 in PBS, Thermo Fisher Scientific, USA). To quantify cells exhibiting lagging chromosomes, 100 anaphase cells were analyzed in each experiment using a Leica DMI6000B fluorescence microscope (Leica, Germany) equipped with a Leica DFC360 FX camera (Leica, Germany) and the Leica LAS AF software (Leica, Germany). Only chromosomes, which were stained with both Hoechst33342 and anti-CENP-C and were clearly separated from the DNA localized at the spindle poles, were considered as lagging chromosomes.

### Detection of W-CIN

To assess time-dependent W-CIN, we analyzed the generation of aneuploidy in single cell clones that were grown for 30 generations in culture. Cells were subjected to chromosome counting analysis from metaphase spreads as described ^10,13^. Briefly, cells were treated for 4 hrs with 2.0 µM of the Eg5 inhibitor Dimethylenanstron (DME) for 4 hrs to accumulate cells in mitosis. Cells were harvested and resuspended in hypotonic solution (60 % ddH_2_O+ 40 % RPMI6140 (PAN-Biotech GmbH, Germany)). After 15 minutes of incubation at room temperature, cells were fixed with ice-cold 75 % methanol + 25 % acetic acid. After fixation, cells were resuspended in 100 % acetic acid and dropped onto pre-cooled wet glass slides. After drying, cells were stained with Giemsa solution (Sigma-Aldrich, Germany). The chromosome number of 50 mitotic cells was quantified using a Zeiss Axioscope FS microscope (Zeiss, Germany) equipped with a Hamamatsu digital camera C4742-95 (Hamamatsu Photonics, Japan) and the Hokawo Launcher 2.1 software (Hamamatsu Photonics, Japan).

### DNA combing assays

DNA combing assays were performed to determine DNA replication fork progression rates and inter-origin distances. Asynchronously growing cells were pre-treated with indicated inhibitors (aphidicolin, ETP-46464, RO-3306, XL-413) for 1 h followed by inhibitor incubation together with 100 μM 5-chloro-2ʹ-deoxyuridine (CldU; Sigma-Aldrich, Germany) and, subsequently, with 100 µM 5-iodo-2ʹ-deoxyuridine (IdU; Sigma-Aldrich, Germany) for 30 min each. Cells were harvested and processed using the FiberPrep DNA extraction kit (Genomic Vision, France). Isolated DNA was immobilized on engraved vinyl silane treated cover slips (Genomic Vision, France) using the Molecular Combing System (Genomic Vision, France). Subsequently, samples were stained with the following antibodies: anti-BrdU (for CldU detection; 1:10, BU1/75 (ICR1), Abcam, UK, cat no ab6326), anti-BrdU (for IdU detection; 1:10, B44, BD Biosciences, USA, cat no 347580), anti-ssDNA (1:5, DSHB, USA, cat no autoanti-ssDNA), secondary antibodies conjugated to Cy5 (1:25, Abcam, UK, cat no ab6565), Cy3.5 (1:25, Abcam, UK, cat no ab6946), and BV480 (1:25, BD Biosciences, USA, cat no 564877). Images were acquired by Genomic Vision’s EasyScan service and samples were analyzed with the FiberStudio web application (Genomic Vision, France). To determine replication fork progression rates, at least 300 labeled unidirectional DNA tracks were analyzed per sample. To analyze inter-origin distances, the distance between two neighboring origins on the same DNA strand was measured. At least 45 inter-origin distances were analyzed per sample.

### Western blotting

Cells were lysed in lysis buffer (50 mM Tris-HCl, pH 7.4, 150 mM NaCl, 5 mM EDTA, 5 mM EGTA, 1 % (v/v) NP-40, 0.1 % (w/v) SDS, 0.1 % (w/v) sodium deoxycholate, phosphatase inhibitor cocktail (25 mM β-glycerophosphate, 50 mM NaF, 5 mM Na_2_MoO_4_, 0.2 mM Na_3_VO_4_, 5 mM EDTA, 0.5 µM microcystin), protease inhibitor cocktail (Roche, Switzerland)). After separation on SDS polyacrylamide gels (7 %, 11 %, or 13 %), proteins were blotted onto nitrocellulose membranes. The following antibodies were used in the indicated dilutions: anti-α-tubulin (1:1000, B-5-1-2, Santa Cruz, USA, cat no sc-23948), anti-β-actin (1:10000, AC-15, Sigma-Aldrich, Germany, cat no A5441), anti-CDC45 (1:1000, D7G6, Cell Signaling Technology, USA, cat no #11881S), anti-CDC7 (1:1000, EPR20337, Abcam, UK, cat no ab229187), anti-MCM2 (1:5000, D7G11, Cell Signaling Technology, USA, cat no #3619S), anti-PSF1 (1:10000, EPR13359, Abcam, UK, cat no ab181112), anti-RIF1 (1:1000, D2F2M, Cell Signaling Technology, USA, cat no #95558), secondary antibodies conjugated to horseradish peroxidase (1:10000, Jackson ImmunoResearch Laboratories, Inc., USA, cat no 115-035-146, 111-035-144). Proteins were detected by enhanced chemiluminescence.

### TCGA molecular and ploidy data

Copy number segment data, gene expression profiles and the ploidy status called by the ABSOLUTE algorithm ^66^ of TCGA primary tumors across 32 cancer types were downloaded from the pan cancer atlas ^67^. Analyzed cancer types included: adrenocortical carcinoma (ACC), bladder urothelial carcinoma (BLCA), breast invasive carcinoma (BRCA), cervical and endocervical cancers (CESC), cholangiocarcinoma (CHOL), colon adenocarcinoma (COAD), lymphoid neoplasm diffuse large B-cell lymphoma (DLBC), esophageal carcinoma (ESCA), glioblastoma multiforme (GBM), head and neck squamous cell carcinoma (HNSC), kidney chromophobe (KICH), kidney cancer (KIPAN), kidney renal clear cell carcinoma (KIRC), kidney renal papillary cell carcinoma (KIRP), acute myeloid leukemia (LAML), brain lower grade glioma (LGG), liver hepatocellular carcinoma (LIHC), lung adenocarcinoma (LUAD), lung squamous cell carcinoma (LUSC), ovarian serous cystadenocarcinoma (OV), pancreatic adenocarcinoma (PAAD), pheochromocytoma and paraganglioma (PCPG), prostate adenocarcinoma (PRAD), rectum adenocarcinoma (READ), sarcoma (SARC), skin cutaneous melanoma (SKCM), stomach adenocarcinoma (STAD), testicular germ cell tumors (TGCT), thyroid carcinoma (THCA), thymoma (THYM), uterine corpus endometrial carcinoma (UCEC), uterine carcinosarcoma (UCS) and uveal melanoma (UVM). A total of 9573 tumor samples, for which copy number segment data, gene expression profiles and the ploidy status data were available, were used for the analysis. The predicted proliferation rates were collected from ^43^.

### Quantifying W-CIN

Copy number and ploidy status were used to compute the weighted genome integrity index (WGII) score for each tumor sample. The WGII score is defined as the average percentage of changed genome relative to the sample ploidy over 22 autosomal chromosomes and ranges from zero to one ^16^.

### Chromosome instability and gene expression association analysis

We first performed gene wise max-min normalization in each cancer type to transform gene expression values to the range between zero. To categorize the tumor samples of a given cancer type as either low or high WGII (W-CIN), we used a k-means based discretization method implemented in the R package *arules* ^68^. To account for cancer type specific effects, we used a meta analysis method implemented in the R package *metafor* ^69^. To estimate the meta-mean difference in gene expression between both WGII groups we used the e*scalc* and *rma* functions in *metafor* with the setting measure=“MD” and method=“FE”. Standard FDR estimates were computed to correct the p-values for multiple testing. Partial correlation coefficients were computed based on the Spearman rank correlation coefficients.

### Chromosome instability and copy number association analysis

The association between chromosome instability and copy number variations (CNVs) were analyzed as for gene expression analysis, replacing gene expression with copy number.

### Gene set enrichment analysis

We used a manually curated list of origin firing genes (*MCM2, MCM3, MCM4, MCM5, MCM6, MCM7, CDC7, DBF4, GINS1, POLD1, POLD2, POLD3, POLE, PCNA, GINS2, GINS3, GINS4, CDC45, CDK1, CCNE1, CCNE2, CDK2, CCNA1, CCNA2, WDHD1, RECQL4, C15orf42, TOPBP1*) and KEGG replication factors (https://www.genome.jp/kegg/) as gene sets to perform gene set enrichment analysis (GSEA) ^70^. All genes were ranked according to their Spearman correlation between expression and WGII and the replication gene or origin firing gene sets were tested for significance enriched at the top of this ranked list.

### Statistical analysis

The GraphPad Prism 5.0 software (GraphPad Software, USA) was used for statistical analysis. Mean values and standard deviation (SD) were calculated. Unpaired two-tailed *t*-tests (SD ≠ 0) or one-sample *t*-tests (SD = 0) were applied to analyze statistical significance. p-values were indicated as: ns (not significant): p≥0.05, *: p<0.05, **: p<0.01, ***: p<0.001, ****: p<0.0001.

## Acknowledgements

We thank Helmut Pospiech and Linda Wordeman for providing plasmids. This work was supported by the FOR2800 funded by the Deutsche Forschungsgemeinschaft (DFG; H.B: FOR2800, sub-project 2 and M.K.: FOR2800, sub-project 3)

The authors declare no competing financial interests.

## Author contributions

Ann-Kathrin Schmidt, Nicolas Böhly, Xiaoxiao Zhang, Benjamin Slusarenko and Magdalena Hennecke designed and performed experiments and analyzed data. Maik Kschischo and Holger Bastians designed the study and analyzed data. All authors contributed to writing the manuscript.

