## Supplemental text for "Dormant replication origin firing links replication stress to whole chromosomal instability in human cancer"

* these authors contributed equally

### Supplemental Figure Legends

**Supplementary Figure S1: Positive correlation of genes involved in replication origin firing with chromosomal instability in human cancer specimens.**

**(a)** Heatmap of Spearman correlation coefficients between the WGII score and the expression of DNA replication and origin firing genes in different cancer types. The genes were ordered by their median correlation over the different cancer types. **(b)** Relationship between copy number variations (CNVs) and WGII scores. The volcano plot shows the mean difference in normalized copy number values in high WGII group versus low WGII group. The meta differences are adjusted for cancer type specific effects in the 32 different tumor types and the p-values are adjusted for multiple testing. Genes of which copy number significantly (FDR $p<0.05$) positively or negatively correlate with W-CIN are marked in blue.

**Supplementary Figure S2: Overexpression of *CDC45* triggers increased origin firing and mitotic errors leading to W-CIN.**

**(a)** Generation of chromosomally stable HCT116 cells with stable *CDC45* overexpression. A representative Western blot shows the expression of endogenous and mCherry-tagged *CDC45* in three independent single cell clones. Single cell clones transfected with empty vector serve as a control. β-actin was used as loading control. Star indicates an unspecific protein band. **(b)** Determination of replication fork progression rates in cells with or without *CDC45* overexpression using DNA combing. Scatter dot plots show values for fork progression rates (mean ± SD, *t*-test). **(c)** Determination of origin firing frequency in cells with or without *CDC45* overexpression. Scatter dot plots show values for inter-origin distances (mean ± SD, *t*-test). **(d)** Determination of mitotic microtubule growth rates in HCT116 cells with or without overexpression of *CDC45* and in the presence or absence of CDC7 inhibition (CDC7i) or Taxol treatment. The indicated single cell clones were treated with 1.0 μM of the CDC7 inhibitor XL-

413 (CDC7i), or with 0.2 nM Taxol for 16 h and microtubule growth rates were determined in mitotic cells. Scatter dot plots show average microtubule growth rates (20 microtubules/cell, n=30 mitotic cells, mean ± SD, *t*-test). **(e)** Quantification of anaphase cells showing lagging chromosomes upon *CDC45* overexpression. The indicated cell clones were treated as in (d) and the proportion of cells with lagging chromosomes was determined. The bar graph shows the quantification of cells with lagging chromosomes (n≥300 anaphase cells, mean ± SD, *t*-test). **(f)** Determination of the proportion of *CDC45* overexpressing cells showing aneuploidy. The indicated single cell clones were grown for 30 generations and chromosome numbers per cell were determined from metaphase spreads. The bar graph shows the proportion of cells with a karyotype deviating from the modal (45 chromosomes in HCT116 cells; n=50 metaphase spreads, *t*-test).

**Supplementary Figure S3: *GINS1* overexpression induces W-CIN.**

**(a)** Determination of the proportion of *GINS1* overexpressing HCT116 cells showing aneuploidy. The indicated single cell clones were grown for 30 generations and chromosome numbers per cell were determined from metaphase spreads. The bar graph shows the proportion of cells with a karyotype deviating from the modal (45 chromosomes in HCT116 cells; n=50 metaphase spreads, *t*-test). **(b)** Depiction of chromosome number deviations in single cell clones shown in (a). Bar graphs show the proportion of cells with the indicated chromosome numbers (n=50 metaphase spreads per clone). Dashed lines represent the average proportion of HCT116 cells with 45 chromosomes in clones transfected with control vector or GINS1 plasmid.

**Supplementary Figure S4: *GINS1* overexpression induces increased microtubule growth rates and chromosomal instability in a CDC7-dependent manner.**

**(a)** Generation of single cell sub-clones overexpressing *GINS1* and continuously treated with 0.5 µM XL-413 (CDC7i) or 0.2 nM Taxol for 30 generations. A representative Western blot shows endogenous and Myc-FLAG-tagged GINS1 protein levels. β-actin serves as loading control. Star indicates an unspecific band. **(b)** Depiction of chromosome number deviations in single cell sub-clones shown in (a). Bar graphs show the proportion of cells with the indicated chromosome numbers (n=50 metaphase spreads per clone). Dashed lines represent the average proportion of HCT116 cells with 45 chromosomes in clones transfected with control vector or GINS1 plasmid. **(c)** Measurements of mitotic microtubule growth rates in single cell sub-clones shown in (a). Scatter dot plots show average microtubule growth rates (20 microtubules/cell, n=10 mitotic cells, mean ± SD, *t*-test).

**Supplementary Figure S5: ATR inhibition increases CDC7-dependent dormant origin firing.**

Determination of origin firing frequencies upon ATR inhibition and concomitant CDK1 or CDC7 inhibition. HCT116 cells were treated with 1.0 µM of the ATR inhibitor ETP-46464 (ATRi), 1.0 µM of the CDK1 inhibitor RO-3306 (CDK1i) or the CDC7 inhibitor XL-413 (CDC7i) for 2 h, pulse-labelled with CldU and IdU and subjected to DNA combing. Scatter dot plots show values for inter-origin distances (mean ± SD, *t*-test).

**Supplementary Figure S6: Knockdown of *CDC7* and CMG components in W-CIN+ colorectal cancer cell lines.**

Representative Western blots show downregulated protein levels of MCM2, CDC7, GINS1, and CDC45 after transfection with siRNAs in **(a)** HCT116, **(b)** SW480, **(c)** SW620, and **(d)** HT29 cells. β-actin was used as loading control. Stars indicate unspecific protein bands.
