## Supplementary figures and images for "Dormant replication origin firing links replication stress to whole chromosomal instability in human cancer"

### Supplemental Figure S1

#### Supplementary Figure 1

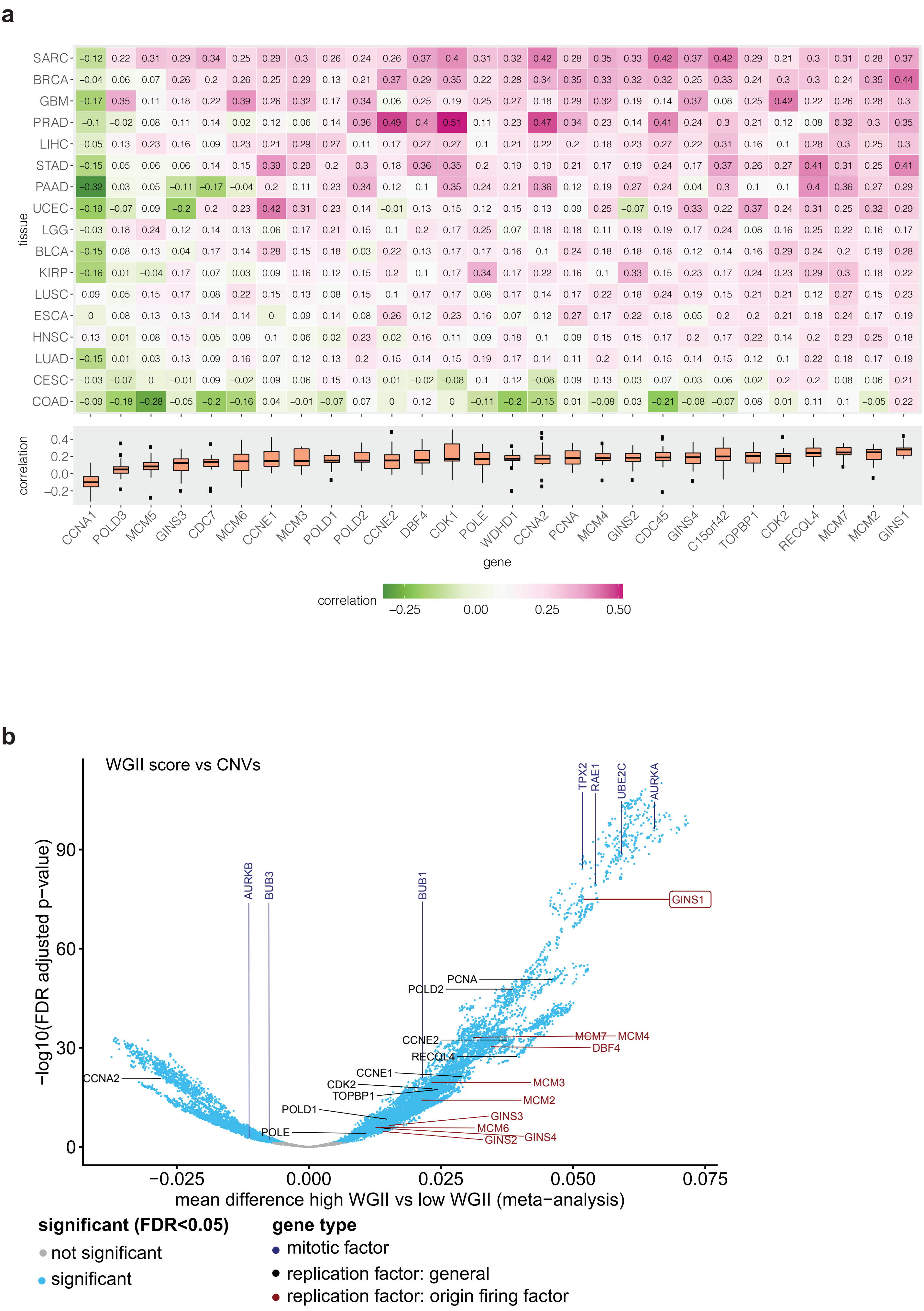

### Supplemental Figure S2

#### Supplementary Figure 2

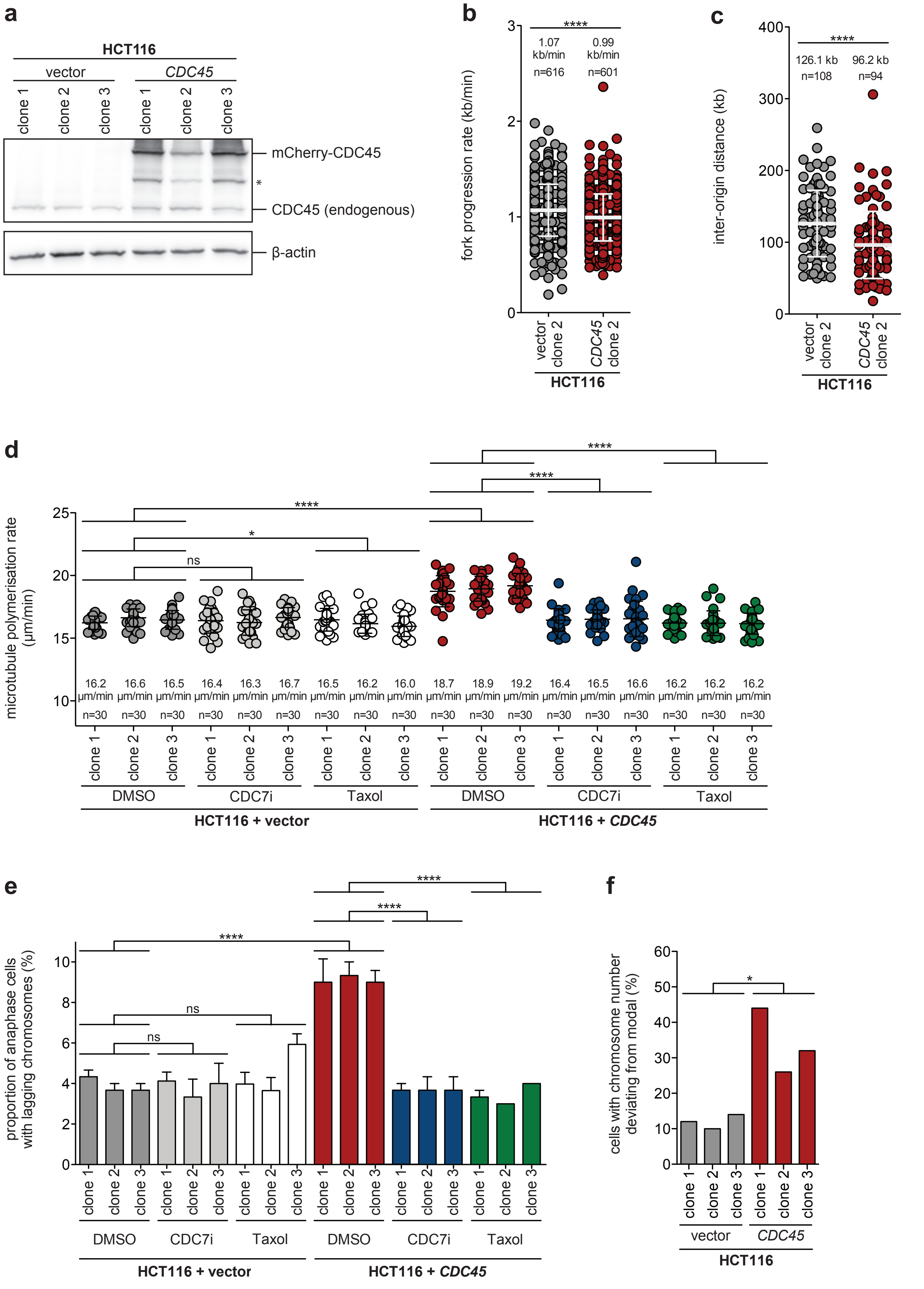

### Supplemental Figure S3

#### Supplementary Figure 3

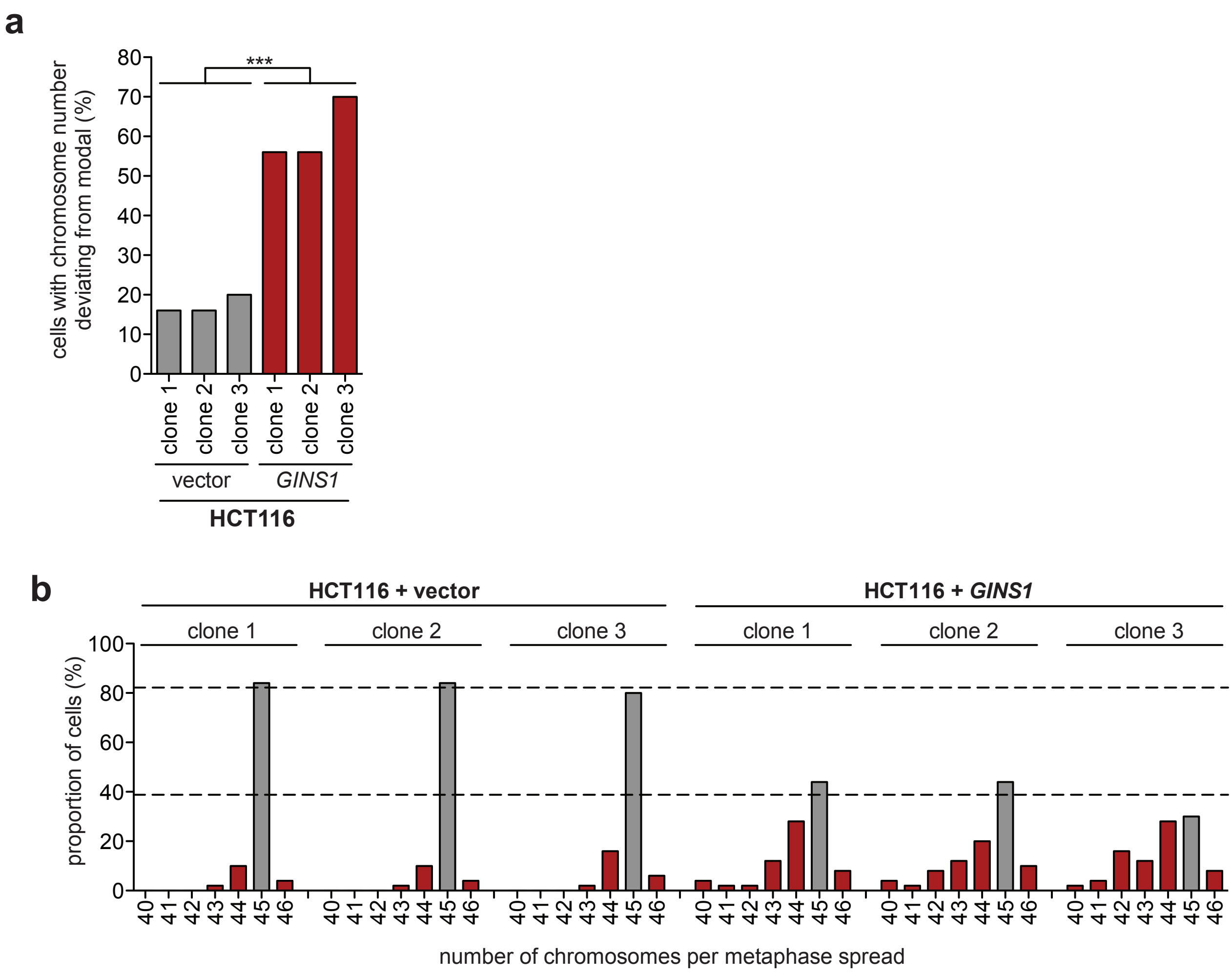

### Supplemental Figure S4

#### Supplementary Figure 4

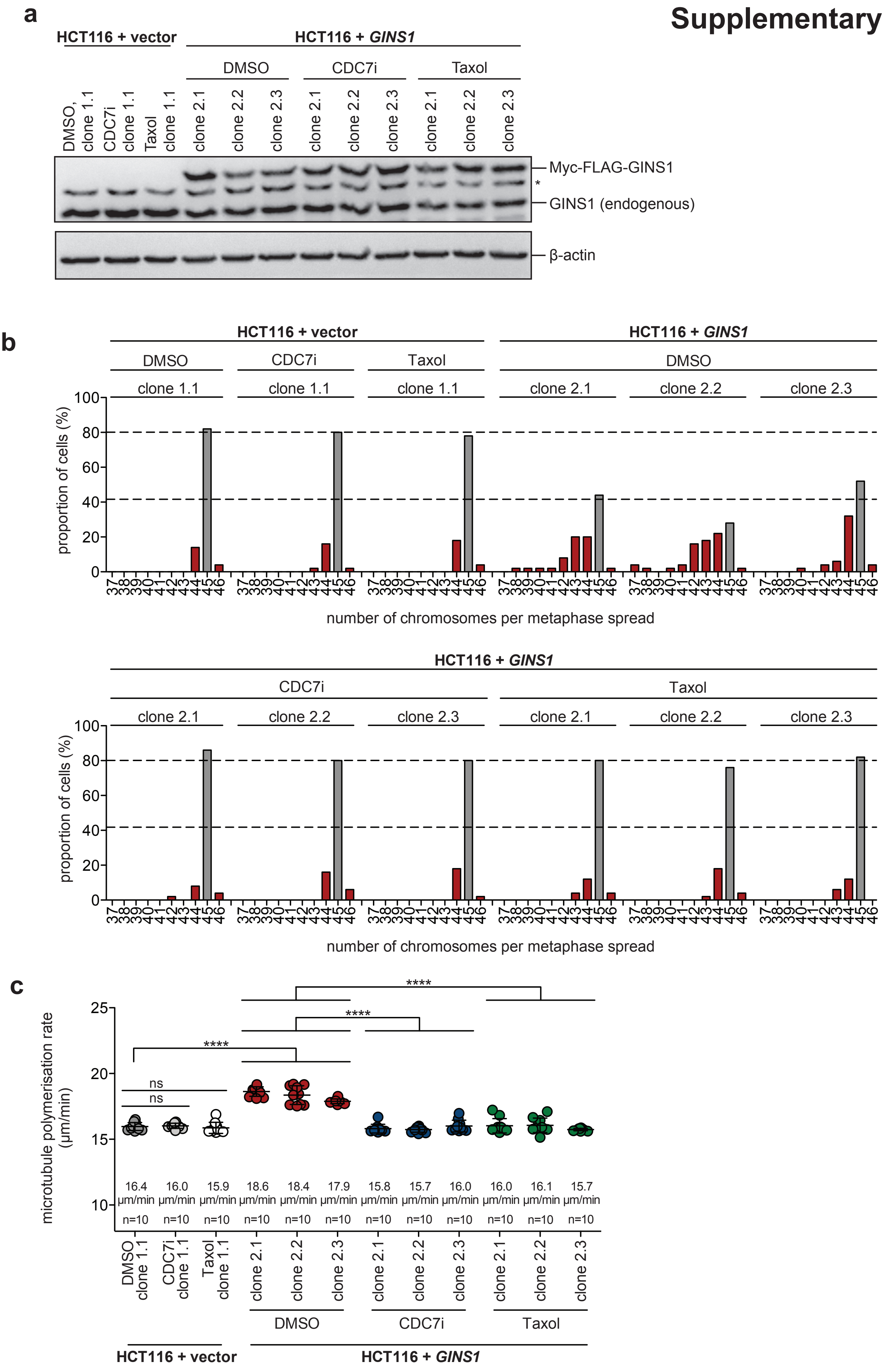

### Supplemental Figure S5

#### Supplementary Figure 5

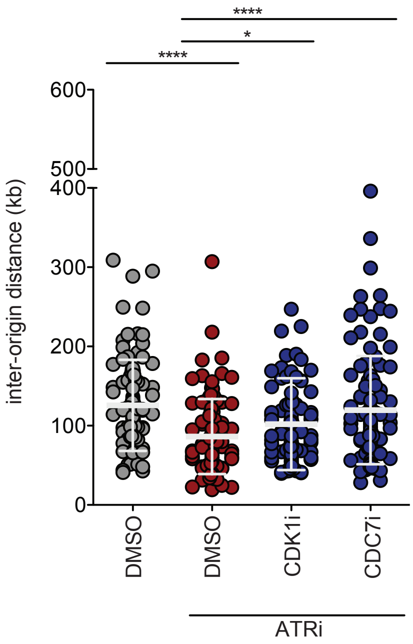

### Supplemental Figure S6

#### Supplementary Figure 6

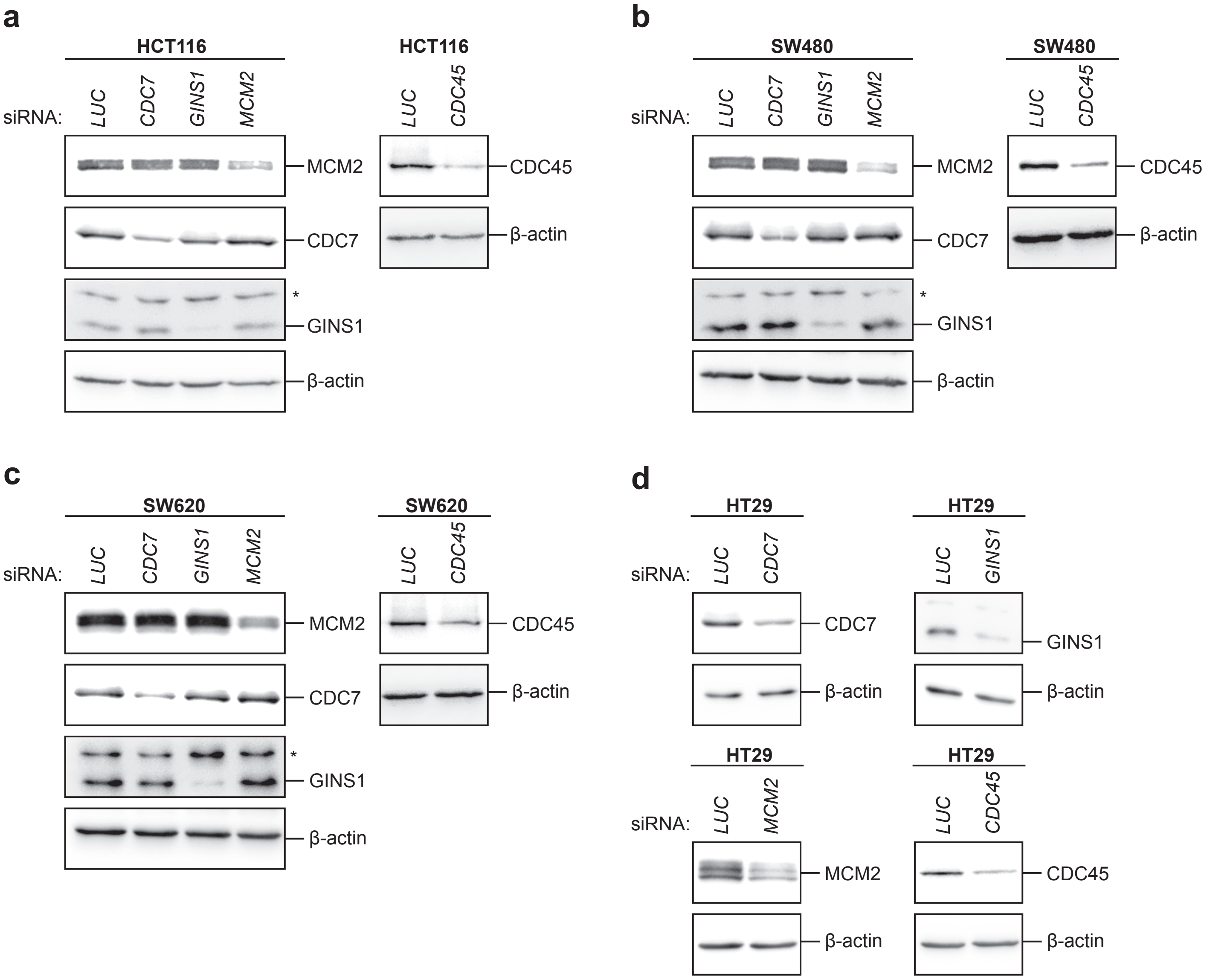
